# Meta-analysis reveals consistent immune response patterns in COVID-19 infected patients at single-cell resolution

**DOI:** 10.1101/2021.01.24.427089

**Authors:** Manik Garg, Xu Li, Pablo Moreno, Irene Papatheodorou, Yuelong Shu, Alvis Brazma, Zhichao Miao

## Abstract

A number of single-cell RNA studies looking at the human immune response to the coronavirus disease 2019 (COVID-19) caused by severe acute respiratory syndrome coronavirus 2 (SARS-CoV-2) have been recently published. However, the number of samples used in each individual study typically is small, moreover the technologies and protocols used in different studies vary, thus somewhat restricting the range of conclusions that can be made with high confidence. To better capture the cellular gene expression changes upon SARS-CoV-2 infection at different levels and stages of disease severity and to minimise the effect of technical artefacts, we performed meta-analysis of data from 9 previously published studies, together comprising 143 human samples, and a set of 16 healthy control samples (10X). In particular, we used generally accepted immune cell markers to discern specific cell subtypes and to look at the changes of the cell proportion over different disease stages and their consistency across the studies. While half of the observations reported in the individual studies can be confirmed across multiple studies, half of the results seem to be less conclusive. In particular, we show that the differentially expressed genes consistently point to upregulation of type I Interferon signal pathway and downregulation of the mitochondrial genes, alongside several other reproducibly consistent changes. We also confirm the presence of expanded B-cell clones in COVID-19 patients, however, no consistent trend in T-cell clonal expansion was observed.

## Introduction

Coronavirus disease 2019 (COVID-19), the global pandemic caused by severe acute respiratory syndrome coronavirus 2 (SARS-CoV-2), is a recognised major threat to the humanity. Thanks to studies from countries around the world, a significant progress in the fields of disease diagnosis, treatment, prevention and control of this disease has been made. However, the pathogenesis of SARS-CoV-2 infection and the immunological characteristics associated with the severity of the disease are still unknown.

To understand the pathology and immune response in COVID-19 patients, a number of single-cell RNA sequencing (scRNA-seq) experiments have been performed on different cell types obtained from human patients^1^–^7^. Studies on diseases caused by influenza and other respiratory viruses have shown that the peripheral immune response plays an important role in the defence against the infections and disease progression^1^. In COVID-19 patients, several pathways have been reported to be regulated, including the CCR1 and CCR5 pathways^5^, the HLA class II and type I interferon pathways^2^, the IL1B pathways and interferon-stimulated genes^4^. However, most studies only focus on pathway analysis in certain cell types. Due to the limited patient sample availability, it is difficult to derive statistically reliable trends in the changes of cell subtype proportions over the disease stages in individual studies. It is still unclear to what extent some of the observations can be generalized and which pathways are consistently regulated.

Here we report results from systematic meta-analysis of 9 single-cell RNA-seq datasets from SARS-CoV-2 infected patient samples and a healthy control dataset. In these datasets, 3 of them have T cell receptor (TCR) sequencing data and 2 have B cell receptor (BCR) sequencing data available. Specifically, we analysed 1) the cell type and cell proportion captured in different experiments to infer the immune response upon SARS-CoV-2 infection; 2) the gene expression regulation and pathway activation in COVID-19 patients; 3) the clonal expansion in T cells and B cells. While we can reproduce 84.21% of the 19 previous conclusions based on small sample size in their original datasets, we could not fully reproduce less than half of these conclusions in other studies. In particular, we show consistent upregulation of type I Interferon signal pathway and downregulation of the mitochondrial genes, as well as demonstrate an absence of consistent T-cell clonal expansion in COVID-19 patients across studies.

## Results

### Meta-analysis reveals common cell types and tissue-specific heterogeneity

#### • Mapping of the disease states

The overview of the datasets is given in **Table 1**, which also gives each dataset a name referred to throughout the text. In total, the datasets that we analysed include 159 experimental samples and 862,354 cells across 9 different disease conditions. Considering the sample stages in different datasets are defined with different names, we map them to standardised terms Healthy, Mild, Moderate, Severe, Post Mild, Convalescent, Late recovery, Asymptomatic and Influenza (**Supplementary Table S1**). Although such retrospective standardisation may introduce a certain level of noise, the mapping of comparable conditions is essential for meta-analysis (**Figure 1**). We have taken care to consider the detailed descriptions of the named conditions as presented in the respective papers and we are confident that for the purposes of our analysis the introduced mappings are correct (see **Supplementary Data 1**).

**Figure 1.**
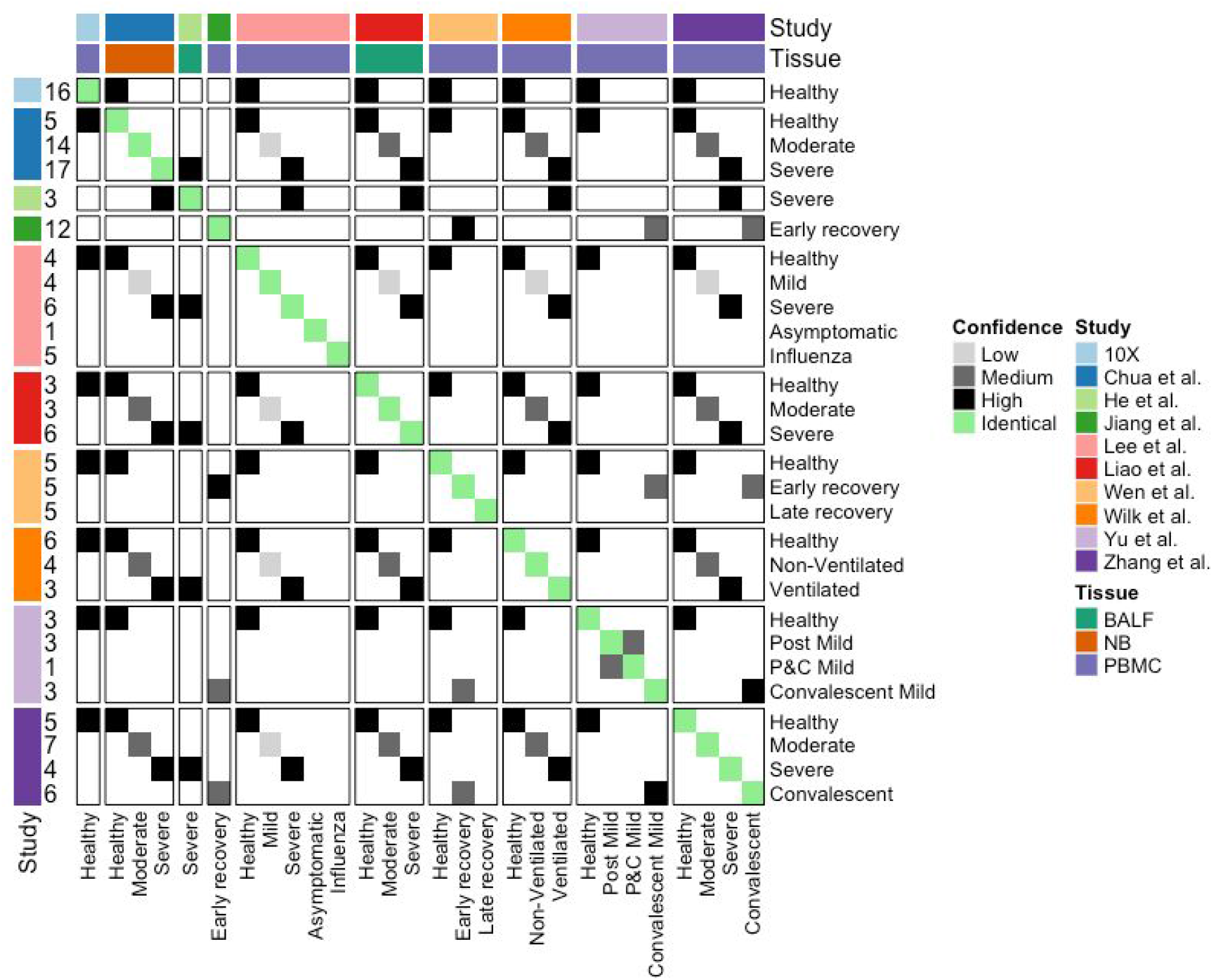
Cross study sample condition comparison. Mapping the conditions in the 10 studies given in Table 1. The number of samples are listed on the left, while the sample tissue types are colored on the top. The similarities between different conditions are colored from white to black (as from no similarity to high similarity). Here, “BALF” denotes samples derived from Bronchoalveolar lavage fluid, “NB” denotes samples derived from nasopharyngeal/bronchial tissue and “PBMC” denotes samples derived from peripheral blood mononuclear cells.

**Table 1.** The scRNA-seq studies of SARS-CoV-2 infected patients samples included in this meta-analysis. The 10 datasets used in this meta-analysis study include 6 datasets (Lee, Wilk, Zhang, Wen, Yu, Jiang) of Peripheral blood mononuclear cells (PBMCs), 2 datasets (Liao and He datasets) of Bronchoalveolar lavage fluid (BALF), 1 dataset of nasopharyngeal and bronchial samples (Chua dataset) and 10X healthy control dataset of PBMCs.

| Study | Accession | Tissue | Technique | 5' or 3' | Samples | Cells | TCR/BCR |
| --- | --- | --- | --- | --- | --- | --- | --- |
| Wen et al. <sup>4</sup> | PRJCA002413 | PBMC | 10X | 5' | 15 | 119,448 | TCR+BCR |
| Zhang et al. <sup>3</sup> | PRJCA002564 | PBMC | 10X | 5' | 22 | 140,588 | TCR+BCR |
| Lee et al. <sup>1</sup> | GSE149689 | PBMC | 10X | 3' | 20 | 58,022 | - |
| Yu et al. <sup>7</sup> | PRJCA002579 | PBMC | 10X | 5' | 10 | 91,742 | - |
| Jiang et al. <sup>24</sup> | NA | PBMC | 10X | NA | 12 | 93,804 | - |
| Wilk et al. <sup>2</sup> | GSE150728 | PBMC | Seq-Well | NA | 13 | 67,923 | - |
| Liao et al. <sup>9</sup> | GSE145926 | BALF | 10X | 3' | 12 | 63,103 | TCR |
| He et al. <sup>6</sup> | GSE147143 | BALF | 10X | 3' | 3 | 10,927 | - |
| Chua et al. <sup>5</sup> | EGAS00001004481 | nasopharyngeal/bronchial | 10X | 3' | 36 | 156,025 | - |
| 10X | NA | PBMC | 10X | 3' | 16 | 60,772 | - |

Visualisation of the combined data using uniform manifold approximation and projection (UMAP) in **Figure 2a** shows strong batch-effect corresponding to different studies. We find the peripheral blood mononuclear cell (PBMC) datasets are more similar to each other and form three large cell clusters, which are the lymphoid lineage, myeloid lineage and the B cells. The Wilk PBMC dataset, which is based on the Seq-Well technique, clusters separately. After the data integration and batch-effect removal with Harmony^8^, the cells from different studies were no longer separated and largely clustered by the underlying biology (**Figure 2b**). Moreover, the healthy samples are also well-mixed, except the epithelial cells, which are mainly captured in the Chua dataset.

**Figure 2.**
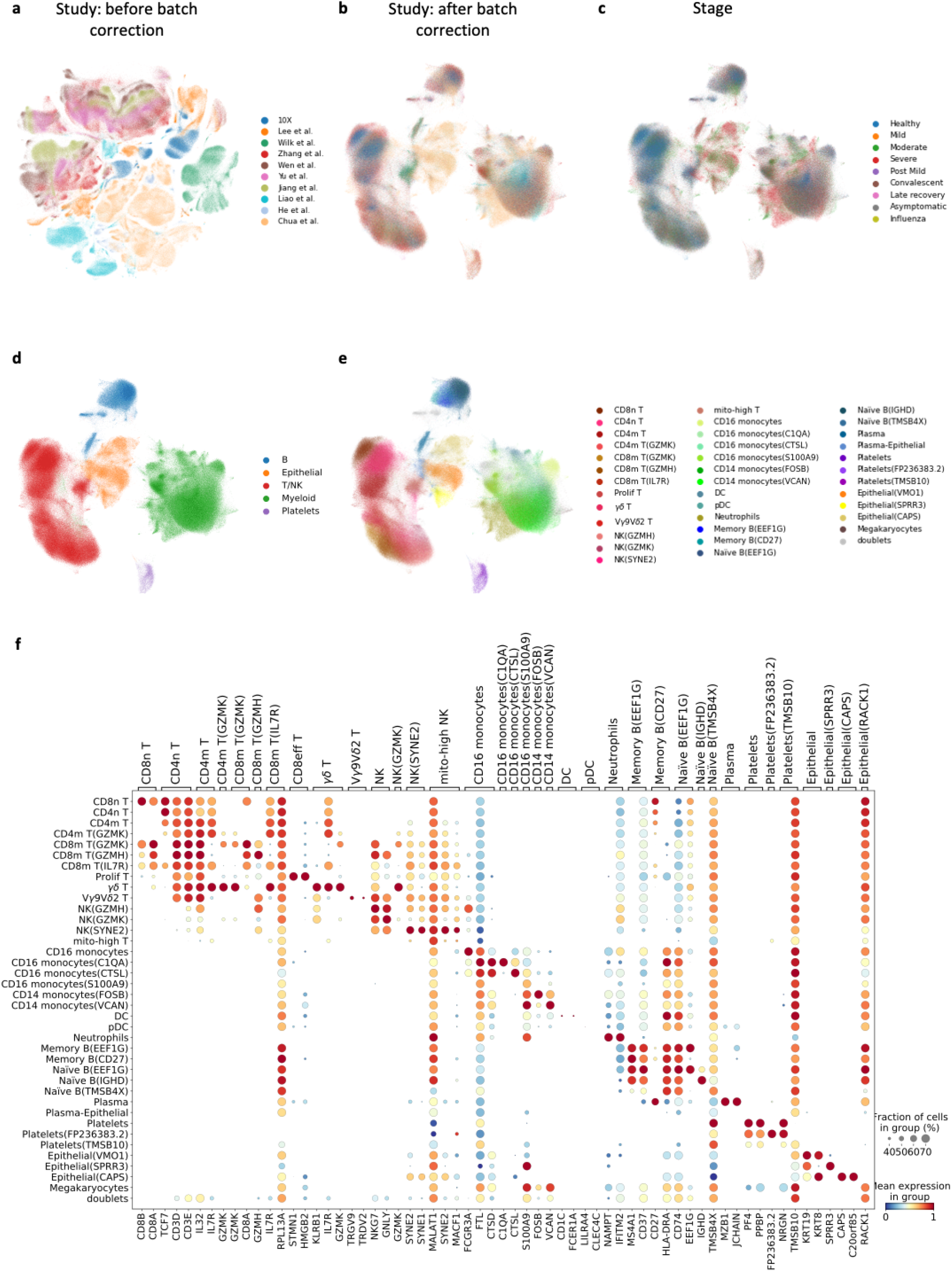
Meta-analysis identifies common and tissue-specific immune cell types. **a)** The UMAP representation of the cells before batch correction, in which the cells are colored by study; **b)** The UMAP representation of the cells after batch correction; **c)** The UMAP representation colored by the disease stage when the sample were taken from the patients; **d)** The UMAP colored by the main cell types; **e)** The description of the cell populations in detail; **f)** The marker genes for discriminating the cell subpopulations.

Similar to previous publication^4,9^, the cells can be classified into 5 major cell populations, **Figure 3**: Lymphoid cells (*CD3D, CD3E, NKG7, GNLY*, Myeloid cells(*CD68, CD14, CD1C, LILRA4*), B cells (*CD79A, MS4A1, MZB1*), Epithelial cells (*KRT8, KRT19*) and Platelets (*PF4, PPBP*). Marker gene expressions are shown in **Supplementary Figure S1**, while previously reported marker gene lists are summarized in **Supplementary Data 2**.

**Figure 3.**
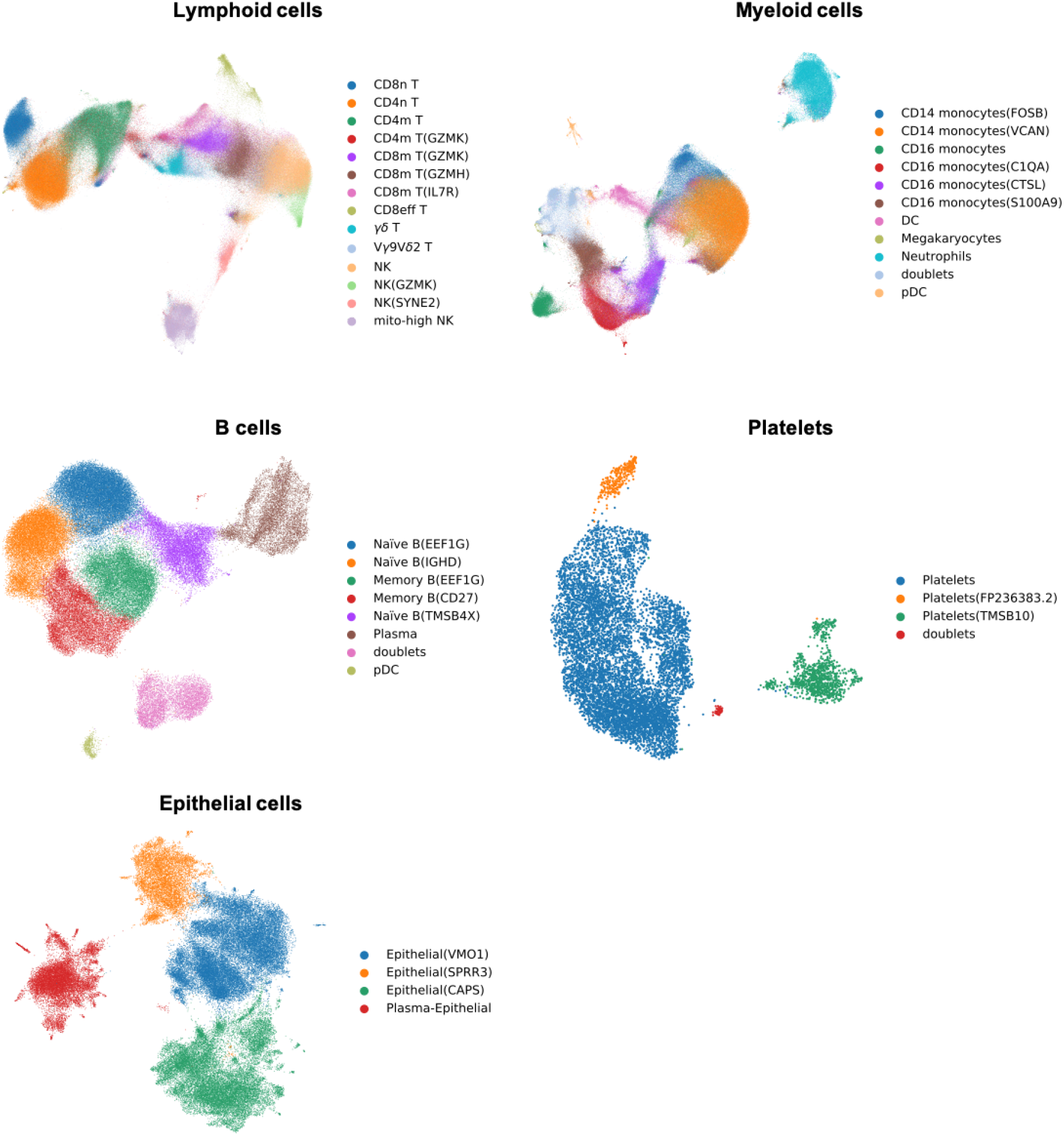
UMAP visualisation of the cell subpopulations in the five main cell types.

Cells in each population are then further divided into subpopulations based on the expression of marker genes and logistic regression reference-based annotation methods (see **Methods)**. The resulting cell clusters were further refined using SCCAF^10^; all the 5 cell populations achieved self-projection accuracies above 92% in the Harmony latent space, **Supplementary Table S2**. The final result includes 37 cell subpopulations excluding doublets.

In the lymphoid cell population, we define 5 CD8^+^ T cell subpopulations, including CD8n T cell, CD8m T(GZMK), CD8m T(GZMH), CD8m T(IL7R) and CD8 effector cells. The CD8n T cells cluster next to the CD4 naive T cells and express *CD8B, CD8A, CD3D, CD3E* and *CCR7*. CD8m T(GZMK), CD8m T(GZMH) and CD8m T(IL7R) are 3 subpopulations of the CD8 T cells and show complementary marker genes (*GZMK, GZMH* and *IL7R*), **Supplementary Figure S2**. *GZMH* is known as a cytotoxic effector T cell marker, while *GZMK* is a transitional effector T cell marker^11^. *GZMH* and *GZMK* not only show good discrimination of the CD8^+^ T cell subpopulations, but also discriminates the NK cell populations, **Supplementary Figure S3**. The CD8eff T cell is a proliferating cell subpopulation and is easily distinguishable.

*CD56 (NCAM1), GNLY* and *NKG7* highlight the NK cells. We further divide the NK cells into 4 subpopulations NK (GZMH^+^), NK(GZMK), NK(SYNE2) and mito-high NK. The mito-high NK cluster has a higher level of percentage of mitochondrial RNA contents and also expresses T cell markers (*CD3D, CD3E*) as well as some red blood cell markers (*HBB, HBA1, HBA2*). Therefore, the mito-high NK cluster is possibly a contaminated subpopulation.

Considering γδ T cells comprise 1%-10% of human PBMCs, it is difficult to distinguish the yō T cell subpopulations within a single dataset. The existing annotations of γδ T cells are not fully consistent: Wilk et al. annotated according to the γδ TCR constant chains encoding genes (*TRGC1, TRGC2* and *TRDC*), while Zhang et al. annotated γδ T cells as TRGV9^+^TRDV2^+^. However, these groups of marker genes highlight two cell subpopulations next to each other, **Supplementary Figure S4**. It is known that Vγ9Vδ2 T cells are the major subset of γδ T cells in human PBMCs^12^. According to the marker knowledge of γδ T cells^12^–^14^, we define the TRGV9^+^TRDV2^+^ subpopulation as Vγ9Vδ2 T cells and other CD161(KLRB1)^+^TRGC1^+^TRGC2^+^ cells as γδ T cells. The γδ T cells also express canonical γδ T cell markers *CCR5, CCR6^15^* as well as *SLC4A10^16^* and *TRAV1-2^17^*, which are known as Mucosal-associated invariant T (MAIT) cell markers.

Although the monocytes cells have been discussed in a previous publication^18^, we are able to refine this type to include 3 CD16^+^ monocytes subpopulations and 2 CD14^+^ monocytes subpopulations. CD16^+^ monocytes express high *FCGR3A (CD16*), while CD16 monocytes(C1QA) not only express the complement proteins (*C1QA, C1QB, C1QC*) but also *HLA-DRA* and *CD63*. The CD16 monocytes(CTSL) cluster expresses *CTSL* and *CD163*, the CD16 monocytes(S100A9) cluster expresses *S100A8* and *S100A9*, **Supplementary Figure S5**. These cell clusters correlate well with the cell populations reported by Schulte-Schrepping et al.^18^. Neutrophils were not annotated in all the reported datasets^34^. However, after annotating neutrophils according to the canonical markers (*FCGR3B* and *CXCR2*), **Supplementary Figure S6**, we find that they exist in all the datasets (but hardly in the healthy controls).

We broadly divided the B cells into naïve B, memory B and plasma cells, where the naive B-cells were further divided into Naïve B(EEF1G), Naïve B(TMSB4X) and Naïve B(IGHD) based on the marker genes’ expression and memory B-cells were further divided into Memory B(EEF1G) and Memory B(CD27).

### Meta-analysis shows consistently decreasing relative proportion of lymphoid cells and increasing proportion of myeloid cells with disease severity

In **Figure 4a**, we show the average cell composition change of the 37 cell subpopulations at different disease stages. The lymphoid cells (T cells and NK cells) decrease from healthy to severe (healthy, mild, moderate, severe), while myeloid cells (CD14 and CD16 monocytes, Neutrophils) relatively increase. In the recovery stages (post mild, convalescent and late recovery), the lymphoid cell proportions are comparable to the proportion at the mild stage but higher than the proportions in moderate and severe stages. And for myeloid cells, the cell proportions are higher than those in the healthy and mild stages. The cell composition of asymptomatic patients is similar to healthy donors. Possibly the asymptomatic patients do not show notable immune responses. Influenza infection data was only found in the Lee dataset and the result shows a very different cell composition from those in COVID-19 infection, where influenza patients have a similar level of lymphoid cells as the moderate COVID-19 patients but more CD14^+^ monocytes.

**Figure 4.**
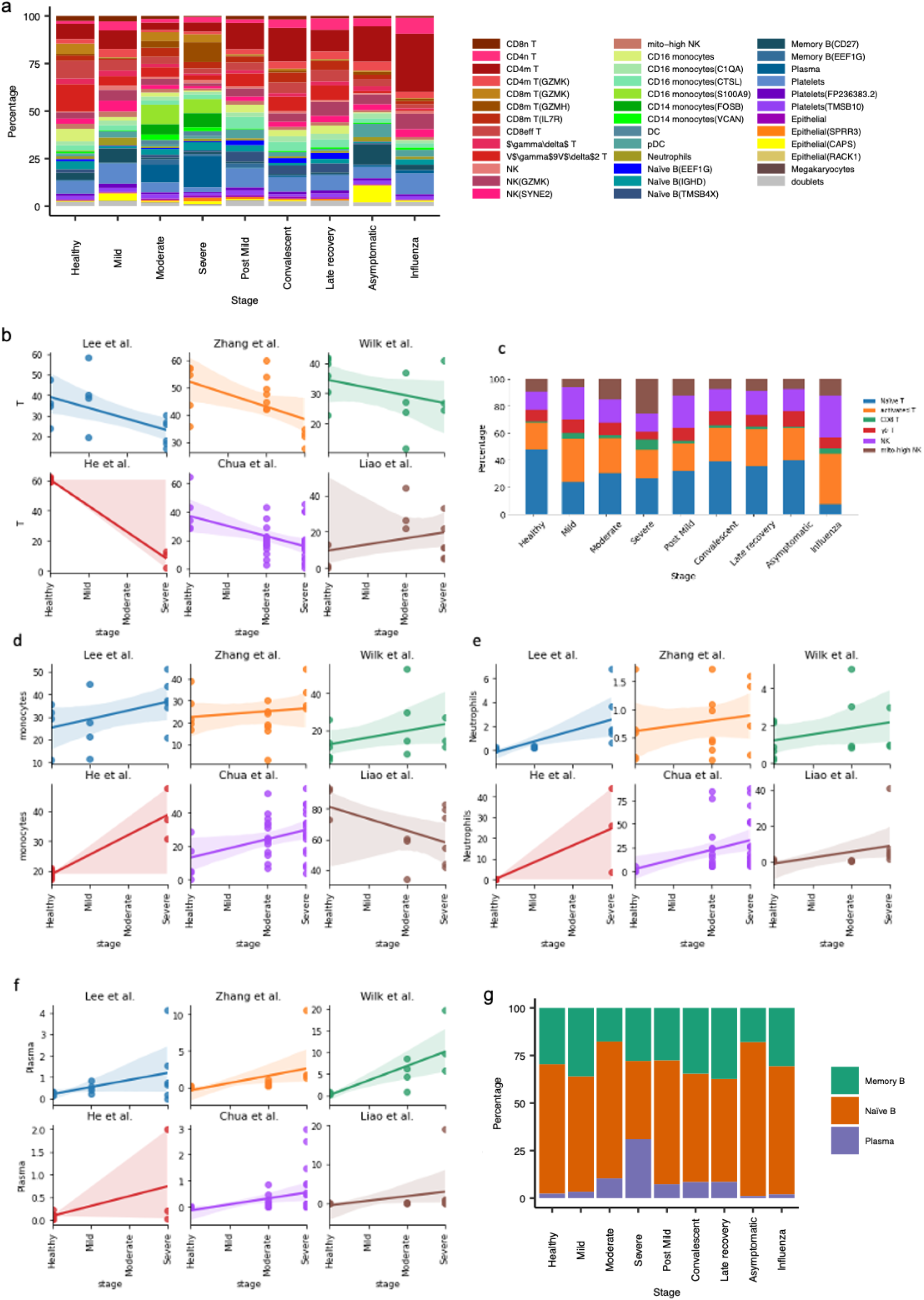
Cell type proportion change upon SARS-CoV-2 infection revealed by multiple studies. **a)** The general distribution of the cell subpopulations at different disease stages; **b),d)-f)** The cell proportion changes upon disease stages for T cells, monocytes, neutrophils and plasma cells, respectively, in different studies. **c)-g)** The general distribution of T-cells/Natural killer (NK) cells and B-cells, respectively, across stages.

To understand the disease process in each dataset separately, 4 datasets (Zhang, Wilk, Chua and Liao) include stages of healthy, moderate and severe, while the Lee dataset includes healthy, mild and severe (here, we keep the mild and moderate as different stages because the samples from patients with mild response show difference in the cell composition from those with moderate infections). Since the 10X reference dataset was used as a healthy reference in the He study^6^, we use the same strategy when analysing this dataset. According to the distribution of these 6 datasets, we find a consistent decreasing trend over the stages from healthy to severe, except for the Liao dataset (**Figure 4b**). Healthy samples of the same tissue type are expected to show similar results. The 4 datasets (Lee, Zhang, Wilk and Chua) show a similar proportion of T cells in healthy donors, which is around 30-40%. But the 10X healthy PBMC reference shows ~60% T cells and the Liao dataset of BALF shows only ~10% T cells. Due to the lack of BALF-derived healthy samples in the He dataset, we cannot confirm whether this is expected in healthy BALF samples or if it corresponds to the general individual-specific variability such as that explored by Wong et al. 2013^19^. However, we can still find a decrease of T cell proportion from moderate to severe stages in the Liao dataset.

The percentages of monocytes and neutrophils relatively increase from healthy to severe stages. Similar to the T cell proportion distribution, the Lee, Zhang, Wilk, Chua and 10X datasets show ~20% monocytes in the healthy samples, **Figure 4d**. In the Liao data this percentage is around 80% and it still shows an increase in monocytes from moderate stage to severe stage. If we use 20% as a reference value for monocytes in healthy controls, in the Liao dataset monocytes further increase from healthy to severe patients. For neutrophils, the 6 datasets consistently illustrate an increasing trend from healthy to severe, though the levels of increase can vary, **Figure 4e**. In the He and Chua dataset, some of the samples may even include 40%-70% neutrophils.

Only a small number of Plasma cells have been detected in the data from healthy donors, but the cell proportion increases in the moderate and severe stages, which is consistent for all the 6 datasets, **Figure 4f**. In contrast, we do not find a consistent cell composition change trend for B cells, **Supplementary Figure S7a**. The B cells increase in the Lee dataset but decrease in the severe samples of the He dataset, while Zhang, Wilk, Chua and Liao datasets do not show a clear trend. Similarly, when looking at B-cells in more granularity, there seems to be a decrease in naïve B-cells in severe patients and a decrease in memory B-cells in moderate and asymptomatic patients compared to healthy controls (**Figure 4g**). However, no consistent trend is observed for naïve B-cells and memory B-cells across multiple datasets (**Supplementary Figure S7b,c**). An increase in naïve B-cell proportion is observed from healthy controls to convalescent (and late recovery patients) in Zhang and Wen datasets (**Supplementary Figure S7d**) but this is contrary to that reported by Wen et al. 2020^4^.

Although dendritic cells take up only a small portion of the population (~2% in healthy PBMCs), they decrease over the disease stages in Lee, Zhang, Chua and Liao datasets (**Supplementary Figure S7e**). Plasmacytoid dendritic cells are even fewer than dendritic cells, the Zhang and Wilk datasets show clear increasing trends over the disease stages (**Supplementary Figure S7f**). But the same conclusion cannot be derived from other datasets (**Supplementary Figure S7f**).

### Robust response to type I interferon and HLA class II downregulation in multiple cell types

To study the response of gene regulatory pathways to COVID infection, we identified differentially expressed (DE) genes and pathways by comparing the mild, moderate and severe patient samples with healthy controls. For each dataset, we first measure the DE genes between moderate samples and healthy controls, and also the DE genes between severe samples and healthy controls, **Figure 5a**. Then, we compare the log fold changes of the genes (False Detection Rate <0.01) between moderate/healthy and severe/healthy. Interestingly, the log fold changes between severe and healthy correlates with the log fold changes between moderate and healthy, which indicates that the genes upregulated and downregulated in moderate and severe samples compared to the healthy controls are similar, **Figure 5b, c**. Moreover, we find the correlation happens in most of the cell types in all studies (including Zhang, Wilk, Liao and Chua). The Lee dataset does not include moderate samples, but includes mild samples, which are different from the moderate ones in cell type composition. It is not surprising that the correlation between mild/healthy and severe/healthy is lower than moderate/healthy and severe/healthy. Nevertheless, this correlation is still positive in all the cell types.

**Figure 5.**
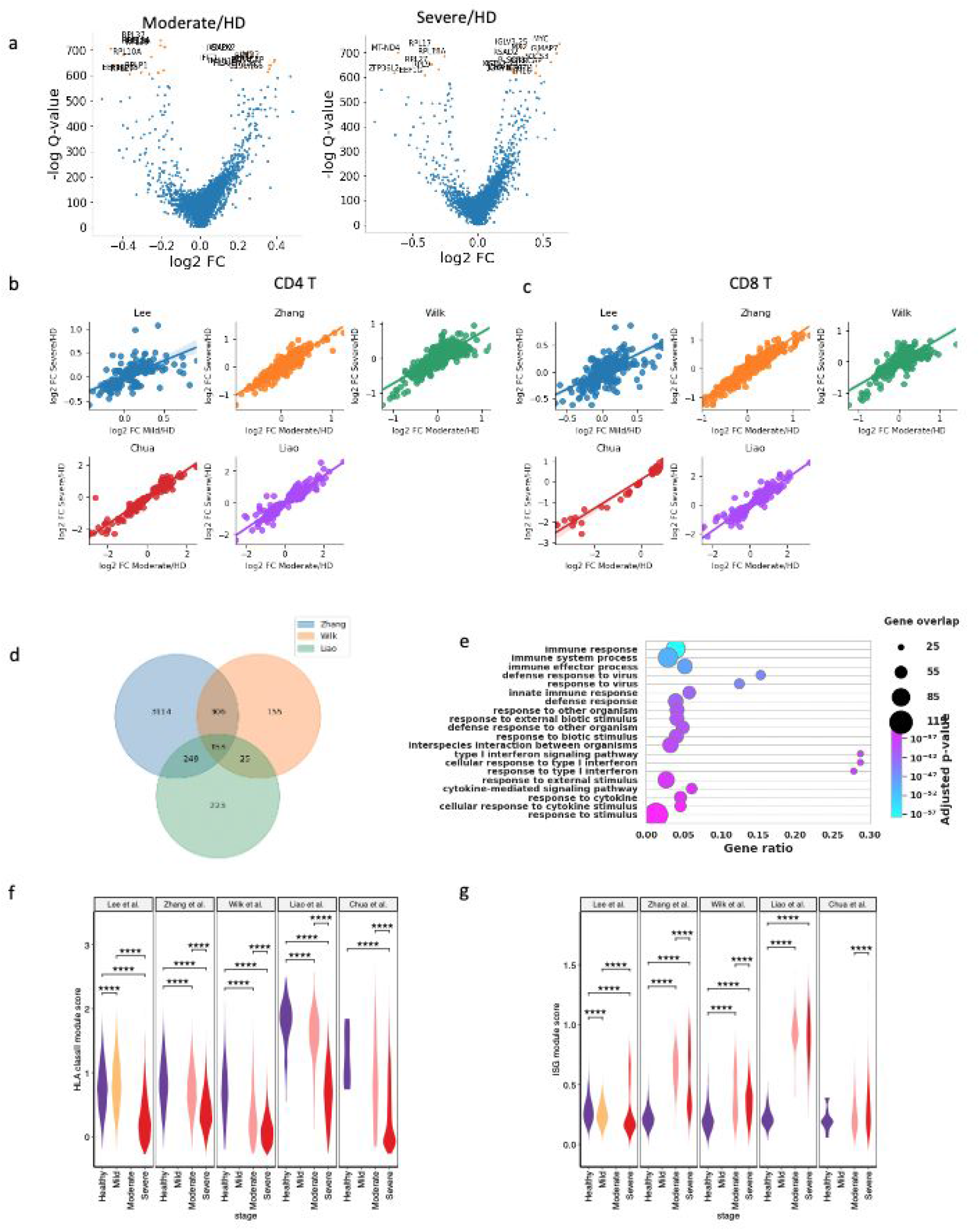
Differentially expressed genes and regulated pathways. **a)** The volcano plots show the differential expressions between: 1) moderate patients and healthy donors; 2) severe patients and healthy donors; **b)** For CD4 T cells, the dot plots show the correlation between the log fold change of moderate/healthy and that of severe/healthy in the five studies; **c)** For CD8 T cells, the dot plots show the correlation between the log fold change of moderate/healthy and that of severe/healthy in the five studies; **d)** The venn plot shows the number of upregulated genes in CD14^+^ monocytes overlapped between the three studies; **e)** The dot plot shows the top upregulated pathways traced from the 153 overlapped genes identified in c). The number of differentially expressed genes (DEGs) that share a particular GO term is represented by the size of the datapoint, the color shows the FDR-corrected p-value, and the x-axis is the ratio of genes in the full DEG set that contain the particular annotation. **f)** Violin plots comparing the HLA class II module score of each CD14^+^ monocytes across studies. All differences were analyzed using two-sided unpaired Wilcoxon rank sum tests with Bonferroni correction and p-values <0.05 are reported. **g)** Violin plots comparing the ISG module score of each CD14^+^ monocytes across studies. All differences were analyzed using two-sided unpaired Wilcoxon rank sum tests with Bonferroni correction and p-values <0.05 are reported.

Furthermore, we identified the genes upregulated in both moderate and severe samples for each study and checked the overlaps between different studies. In CD14 monocytes, 3822, 639 and 650 genes are upregulated in the Zhang, Wilk and Liao datasets respectively. Of these, 153 genes are consistently upregulated by all the studies, **Figure 5d**. Pathway analysis of these 153 genes highlight the gene ontology terms: immune response, response to virus and type I interferon signalling pathway. Although the upregulation of type I interferon signalling pathway in monocytes has been reported by Wilk et al. 2020^2^, the upregulation mainly happens in CD14 monocytes and not in CD16 monocytes. We find the upregulation of type I interferon signalling pathway in many of the cell types, including T cells (CD4 T, CD8 T and γδ T), NK cells, DC, pDC, B cells, Plasma cells and Neutrophils (**Supplementary Figure S8**). Thus, the upregulation of type I interferon signalling pathway seems to be consistent for many cell types. Specifically, we find the upregulation of *IFITM3, IFIT3, IFI44, IFI44L, IFI16, IFI6, LY6E, ISG15, ISG20, OAS1, OAS2, OAS3* and *OASL*.

Similarly, genes related to SRP-dependent co-translational protein targeting to membrane as well as the mitochondrial proteins are consistently downregulated in immune cell types. In NK cells and γδ T cells, 16 and 17 genes are downregulated in all the three datasets (Zhang, Wilk, Liao) respectively. These genes include mitochondrial genes (*MT-ATP6, MT-CO1, MT-CO2, MT-CO3, MT-CYB, MT-ND4, MT-ND5, MT-ND3, MT-ND1, MT-ND2) KLRB1* and *UBA52*.

Additionally, the human leukocyte antigen (HLA) class II genes have been reported to be downregulated in monocytes of severe COVID-19 patients^2^. We found that in CD14 monocytes, HLA class II module scores (see **Methods**) were significantly lower (Bonferroni corrected p-value < 0.05) in severe COVID-19 patients than healthy controls across all the three PBMC datasets (Lee, Zhang and Wilk), as well as in the Liao dataset and the Chua dataset (**Figure 5f**). This observation was also consistent in CD16 monocytes, dendritic cells (DCs) and plasmacytoid dendritic cells (pDCs) across all the five datasets but not in NK cells, naive B-cells or memory B-cells (**Supplementary Figure S9**).

The interferon-stimulated genes (ISGs) have been reported to be heterogeneously upregulated in CD14 monocytes in COVID-19 (both moderate and severe)^2^. We found the result is reproducible in Wilk, Zhang and Liao datasets. However, the result in the Lee dataset demonstrates a downregulation and the difference in Chua dataset was not statistically significant (**Figure 5g**). Therefore, we may not confirm a definite result for the ISG regulation in COVID-19 patients.

### Meta-analysis did not provide validation of the developing neutrophil population in severe COVID-19 patients

Although we were able to reproduce the developing neutrophil population described by Wilk et al. 2020^2^ in their own dataset (**Supplementary Figure S10d**), we could not validate it in the severe COVID-19 patients from other datasets (**Supplementary Figure S11**). In fact, we noticed that in Wilk et al. 2020 too these developing neutrophil markers were present in the cell subpopulation annotated as “Epithelial (RACK1)” (**Supplementary Figure S10a,d**). When subsetting this cell subpopulation only to the severe patients of Wilk dataset, we noticed that it had marker genes for red blood cells, monocytes, developing neutrophils, B-cells and 2 other clusters, thereby representing a broad set of cells (**Supplementary Figure S10b-c**). To make sure that we didn’t miss the developing neutrophil population in the severe COVID-19 patients from other datasets during the sub-setting process, we first checked all the developing neutrophil marker genes’ expression for all the cell-types and then only for plasma cells and epithelial (RACK1) cells. We did not observe the developing neutrophil population in either case (**Supplementary Figure S11**). This may partly be explained by the low number of plasma and epithelial (RACK1) cells captured by other datasets compared to Wilk dataset (n=2862 cells). Also, Seq-Well platform for scRNA-seq used in Wilk dataset compared to 10X used in other studies might have an effect on top of the underlying patient-specific variability contributed by factors such as age, sex, ethnicity, genetics, received immunomodulatory drugs and lifestyle. Further, the datasets Liao et al. 2020^9^ and Chua et al. 2020^5^ were derived from BALF and nasopharyngeal/bronchial tissues, respectively, compared to others derived from PBMC.

### Meta-analysis is inconclusive in regards to a consistent trend in T-cell clonal expansion across COVID-19 stages

For the current section, we analyzed the T-cell receptor (TCR) data from Wen et al. 2020^4^, Zhang et al. 2020^3^ and Liao et al. 2020^9^ where the samples from the first two studies were derived from PBMC while those from the third one were derived from BALF. Among these, the conditions healthy and convalescent were present in both the PBMC datasets, while the moderate and severe stages were present in Zhang et al. 2020^3^ and Liao et al. 2020^9^ TCR datasets only.

We first pooled T-cell and NK cell data from all the studies together and visualized them using UMAP (**Figure 3**: “Lymphoid cells” and **Figure 6a**). As expected, NK-cells mostly had no corresponding TCR data. For the cells with TCR data, we observed clonal expansion of CD8m T-cells, particularly the subpopulations CD8m T(GZMH) and CD8m T(GZMK) and Vy9V□2 T-cells (**Figure 6a**). Their clonal expansion did not appear to be stage-specific (**Figure 6a**). However, we observed an over-representation of CD8^+^ effector T-cells in severe patients. These observations were confirmed in the bar plots where samples from all the datasets at a specific stage were pooled in together and normalized according to the total number of cells at each stage (**Figure 6b**). This is also consistent with Zhang et al. 2020^20^.

**Figure 6.**
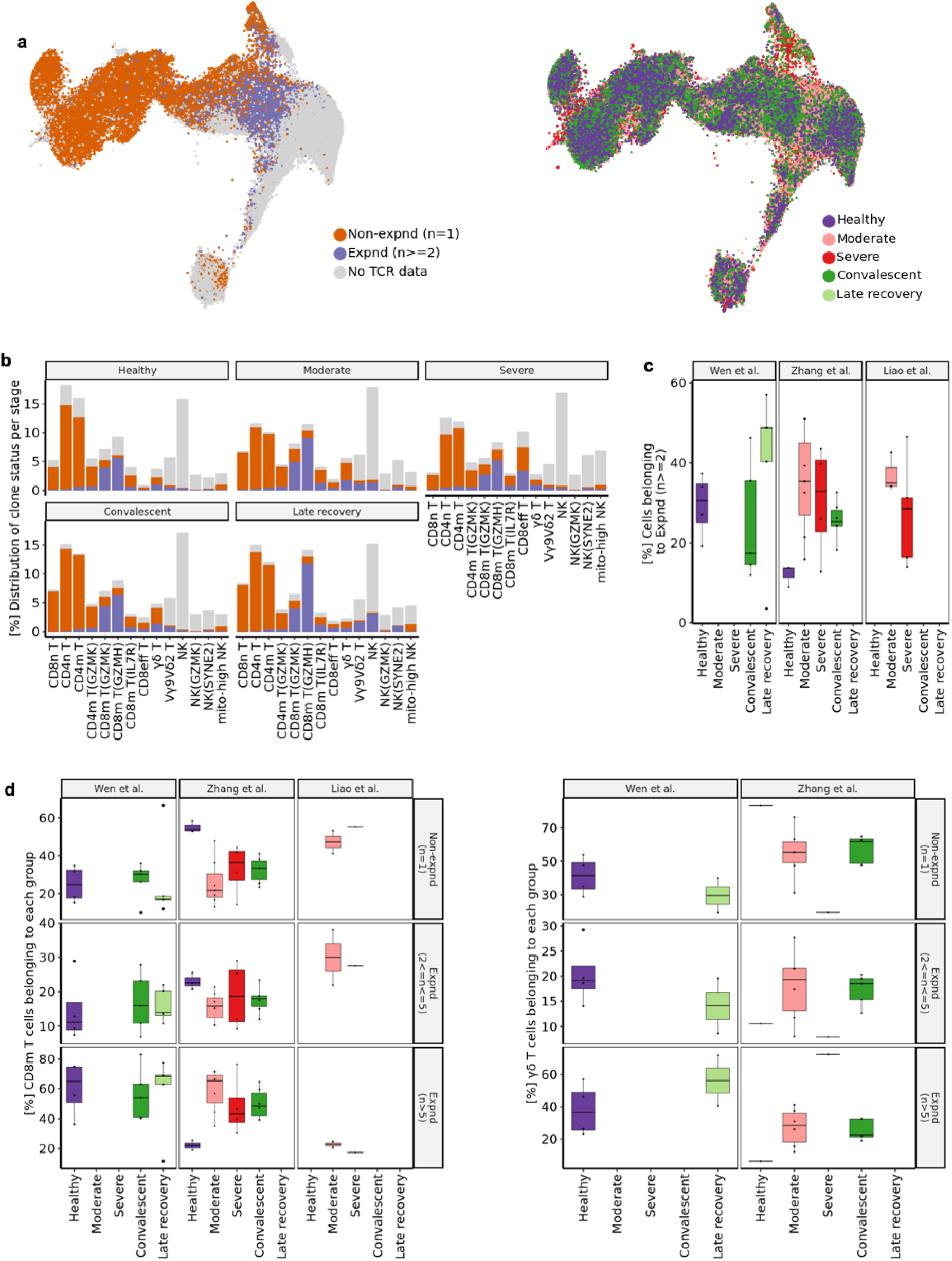
Meta-analysis yields inconclusive results regarding the CD8^+^ T-cell clone expansion in COVID-19 patients compared to healthy. **a)** UMAP of T cells and NK cells derived from Wen et al. 2020, Zhang et al. 2020 and Liao et al. 2020 (BALF). Colors indicate the clusters of TCR detection and cells belonging to expanded (Expnd) vs non-expanded (Non-expnd) clonotypes (left) and the stage of the patients (right). Here, n denotes the number of clones in each clonotype belonging to that group. **b)** Bar plots showing the percentage of cells at each stage belonging to the specific T-cell subpopulation as well as clone status. Here, grey color denotes no TCR data, orange color denotes non-expanded clone status (Non-expnd) and blue color denotes expanded clone status (Expnd). **c)** Box plots comparing the percentage of all the T-cells belonging to expanded clonotypes (Expnd (n>=2)) at each stage across studies. Colors denote the stage of the patient. All differences were analyzed using two-sided unpaired Wilcoxon rank sum tests with Bonferroni correction and p-values <0.05 are reported. **d)** Box plots comparing the percentage of CD8^+^ memory T-cells (left) and γ□ T-cells (right) belonging to each group of clonotypes at each stage across studies. Here, the three groups of clonotypes are non-expanded clonotypes (Non-expnd (n=1)), expanded clonotypes with greater than 2 and less than 5 clones (Expnd (2<=n<=5)) and expanded clonotypes with greater than 5 clones (Expnd (n>5)). Colors denote the stage of the patient. All differences were analyzed using two-sided unpaired Wilcoxon rank sum tests with Bonferroni correction and p-values <0.05 are reported.

To further check if these observations were consistent across studies, we analyzed the overall clonal expansion trend of T-cells. Contrary to previously published results^3,4^, we didn’t observe a consistent trend of increased clonal expansion of T-cells in COVID-19 patients compared to healthy controls (**Figure 6c**). Although the moderate and severe patients of Zhang et al. 2020^3^ and Liao et al. 2020^9^ seemed to share a similar percentage range of clonally expanded T-cells, Liao et al. 2020^9^ lacked the TCR data corresponding to healthy controls from BALF tissue, making it difficult to draw reliable conclusions. Also, the healthy controls from Zhang et al. 2020^3^ seemed to be different from the healthy controls from Wen et al. 2020^4^ in terms of percentage of clonally expanded T-cells, thereby making it difficult to find a decrease in convalescent patients compared to healthy controls as reported by Wen et al. 2020^4^.

Zhang et al. 2020^20^ showed that the overall increased TCR clonal expansion was observed in the severe convalescent COVID-19 patients, compared to healthy control, in the top 6 clonotypes ratio analysis. However, in all datasets, the differences between individuals were found to be large, and in some datasets greater than the differences between groups (**Figure 6c**)^20^. This seems to indicate that there are multiple factors that may affect TCR clonal expansion.

We repeated the analysis on CD8^+^ T-cell subpopulations showing clonal expansion in **Figure 6a** (and **Figure 6b**) to check if they share a common trend across stages. As shown in **Figure 6d**, the results were inconclusive for memory CD8^+^ T-cells but increased clonal expansion of γ□ T-cells, with more than 5 clones in each clonotype (group ‘Expnd (n>5)’) was observed in recovered patients compared to healthy across both the PBMC datasets. Further, due to the presence of less than 100 CD8^+^ effector T-cells in most of the samples, we were unable to analyze this subpopulation in greater detail (**Supplementary Figure S12a-b**). We also observed that all the samples from the Liao et al. 2020^9^ TCR dataset had less than 100 cells each for most of the T-cell subpopulations (**Supplementary Figure S12a**), thereby leaving us with only one dataset, Zhang et al. 2020^3^, with samples from severe and moderate stages. Regarding the remaining T-cell subpopulations, naive CD8^+^ T-cell showed increased clonal expansion in severe patients in Zhang et al. 2020^3^ compared to healthy control (clonally expanded clonotypes with 2 to 5 clones each; group ‘Expnd (2<=n<=5)’) and similar clonal expansion status in convalescent patients compared to healthy controls in Zhang et al. 2020^3^ (**Supplementary Figure S12c**). Whereas for naive CD4^+^ T-cells, the convalescent stage appeared to be similar to healthy across both the PBMC datasets (**Supplementary Figure S12d**). Similar to memory CD8^+^ T-cells, we didn’t find any consistent trend in memory CD4^+^ T-cells (**Supplementary Figure S12e**).

### Meta-analysis of B-cell receptor data confirms the presence of clonally expanded B-cells in COVID-19 patients across multiple datasets

The B-cell receptor (BCR) data was collected from two PBMC studies: Wen et al. 2020^4^ and Zhang et al. 2020^3^ out of which the stages healthy and convalescent were present in both the datasets. The initial visualization of B-cell subpopulations with or without BCR data using UMAP showed that the BCR data was available for almost all the B-cell subpopulations (**Figure 3**: “B-cells” and **Figure 7a**). It also showed the presence of expanded Plasma cells in mostly COVID-19 patients (**Figure 7a**). This observation was confirmed when cells from all the studies at each stage were taken together and normalized by the total number of cells present at each stage (**Supplementary Figure S13a**). Increased clonal expansion of Naïve B(EEF1G) and Memory B(EEF1G) cell subpopulations was also observed in severe COVID-19 patients compared to other B-cell subpopulations in severe patients (**Supplementary Figure S13a**).

**Figure 7.**
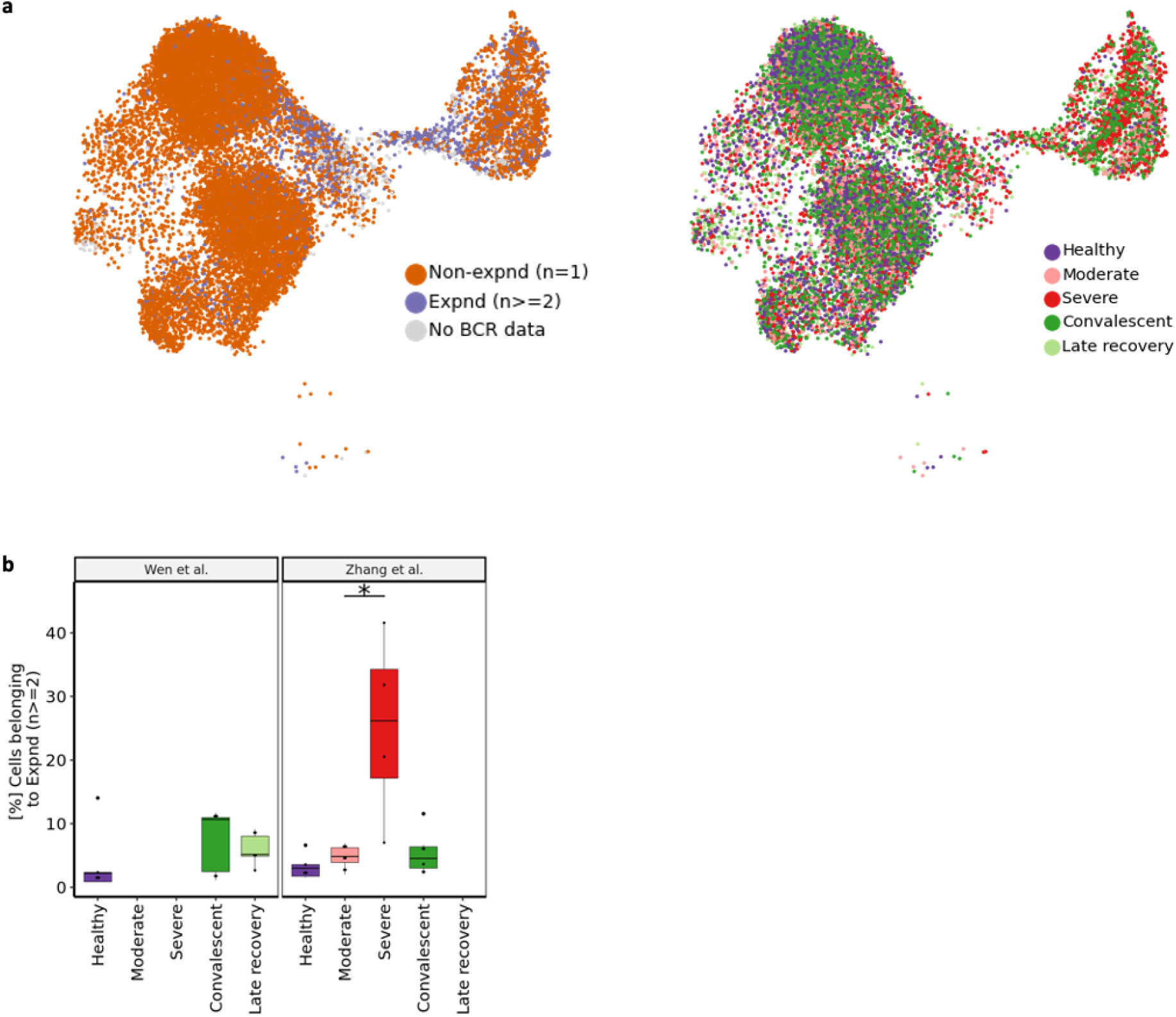
Meta-analysis confirms the presence of expanded B-cell clones in COVID-19 patients. **a)** UMAP of B cells derived from Wen et al. 2020 and Zhang et al. 2020. Colors indicate the clusters of BCR detection and cells belonging to expanded (Expnd) vs non-expanded (Non-expnd) clonotypes (left) and the stage of the patients (right). Here, n denotes the number of clones in each clonotype belonging to that group. **b)** Box plots comparing the percentage of all the B-cells belonging to expanded clonotypes (Expnd (n>=2)) at each stage across studies. Colors denote the stage of the patient. All differences were analyzed using two-sided unpaired Wilcoxon rank sum tests with Bonferroni correction and p-values <0.05 are reported.

We further analyzed whether the previously reported observations^3,4^ of increased B-cell clonal expansion in COVID-19 patients compared to healthy controls can be confirmed and found that this is indeed consistent (**Figure 7b**). For closer examination of clonal expansion of specific B-cell subpopulations in COVID-19 patients (**Figure 3**: “B-cells”), we grouped them into three major groups to have a comparable number of cells in each sample: namely memory B-cells, naïve B-cells and plasma cells. It was observed that there were less than 100 plasma cells with corresponding BCR data in most of the samples to draw any reliable conclusions (**Supplementary Figure S13b**). In case of naïve B-cells, we observed slightly higher clonal expansion in convalescent COVID-19 patients compared to healthy controls in both the studies (clonally expanded clonotypes with 2 to 5 clones each; group ‘Expnd (2<=n<=5)’; **Supplementary Figure S13c**), but this difference was not statistically significant. We also observed increased clonal expansion of naïve B-cells in moderate and severe COVID-19 patients compared to healthy controls in both the clonally expanded groups (‘Expnd (2<=n<=5)’ and ‘Expnd (n>5); Supplementary Figure S**13c**) but as the Wen dataset lacks samples from these stages, this observation is not applicable to the Wen dataset. In case of memory B-cells we didn’t observe any consistent trend in clonal expansion status across stages (**Supplementary Figure S13d**)

## Discussion

Understanding the immunological landscape of SARS-COV2 is an important part of molecular biology research relevant to fighting the pandemic. Previous single cell studies have tried to contribute to the knowledge of COVID-19 infection across multiple disease stages, however, these studies suffer from limited sample size and lack standardization in data collection, pre-processing, cell-type annotation and analysis. In this study, we have tried to identify which of the previously published conclusions are reproducible across multiple datasets and which are study specific when these datasets are analyzed in a standardised way (summarized in **Tables 1 and 2**).

By remapping the reads, we were able to identify certain cell populations that were missed out by the original studies. For example, we observed neutrophil population in the Zhang and the Wen datasets, not reported by the original studies. Further, all nine COVID-19 scRNA-seq datasets analyzed in this study had samples collected from patients at different stages, thereby making it difficult to perform homogeneous comparison across all studies. An ideal way would have been to regroup the samples according to the stages defined on a standardized scale (e.g. the WHO scale^21^) but due to the lack of this metric in the present datasets, we had to make various assumptions, which we verified by considering the available detail in the information provided in original publications and in some cases by contacting the authors. In particular, we assumed that the healthy, severe, early recovery (convalescent) and asymptomatic stages should be comparable. We also checked whether samples at moderate stage from other datasets could be merged with samples at mild stage from Lee et al. 2020^1^ dataset, but these two stages exhibited different cell-proportion distributions (Figure 4) and differentially expressed genes and were, therefore, kept as separate. Presence of more BCR data, longitudinal samples with a detailed history of received therapeutic interventions, particularly those with immunomodulatory effects such as Azithromycin^22^ given to severe patients in the Wilk dataset^2^, additional clinical data on pre-existing conditions and ethnicities^19^, comparison with asymptomatic patients and presence of other omics modalities^23^–^27^ would have helped to strengthen the current analysis.

Bar the mentioned uncertainties, which may explain some of the unconfirmed original results across individual studies, this study provides a direct comparison of multiple single cell COVID-19 studies published to date. As shown in **Table 2**, we can reproduce 16 out of 19 (84.21%) previously published results we considered in the original datasets and can only reproduce 8 out of 19 (42.11%) previously published results in other datasets.

**Table 2.** Summary of reproducible results across multiple single-cell COVID-19 studies analyzed in this meta-analysis. TCR: T-cell receptor; TNF: tumor necrosis factor; IL: interleukin; HLA: human leukocyte antigen; CD8eff T-cell: CD8^+^ effector T-cell

| Study | Key finding(s) in COVID-19 patients compared to healthy controls | Reproduced in the original dataset | Consistently reproduced in all other dataset(s) considered for comparison | Potential reason(s) in case of non-reproducibility |
| --- | --- | --- | --- | --- |
| Wen et al. <sup>4</sup> | Decreased CD4<sup>+</sup> and CD8<sup>+</sup> T-cells (data not shown) | Yes | Yes |  |
|  | Increased CD14<sup>+</sup> monocytes (Supp Fig. S7h) | Yes | No | Patient-specific variability in fighting COVID-19 infection; lack of comparable stages in other datasets |
|  | Increased B-cell clonal expansion (Fig. 7b) | Yes | Yes |  |
|  | Increased Plasma cells (Fig 4f, Supp Fig. S7h) | Yes | Yes |  |
|  | Decreased naïve B-cells (Supp Fig. S7c, S7d) | No | No | Patient-specific variability in fighting COVID-19 infection |
| Zhang et al. <sup>3</sup> | IFN-α response upregulation (Figs 5d, 5e and Supp. Fig. S8) | Yes | Yes | |
|  | Increased T-cell clonal expansion (Fig. 6c) | Yes | No | Lack of comparable stages across multiple datasets having TCR data |
|  | Increased CD8eff T-cell clonal expansion (Supp Fig. S12b) | Yes | Yes |  |
|  | Increased Plasma cells (Fig. 4f) | Yes | Yes |  |
|  | Decreased memory B-cells (Supp Fig. S7b) | Yes | No | Patient-specific variability in fighting COVID-19 infection |
| Lee et al. <sup>1</sup> | TNF/IL-1β driven inflammatory response upregulation (data not shown) | No | No | Not all the cell types are consistent. In the Lee dataset, TNF and IL1B show higher expression in moderate samples but not too high in severe samples. In case of other datasets, different studies are different. e.g., Wilk T cells do not express significant TNF/IL1B. |
| | Co-existence of type I IFN response with TNF/IL-1β-driven inflammation in severe COVID-19 patients (data not shown) | Yes, but not sharp contrast with milder | No | Patient-specific variability in fighting COVID-19 infection |
| Wilk et al. <sup>2</sup> | Developing neutrophil population from plasmablasts in severe COVID-19 patients (Supp Figs. S10, S11) | Yes | No | Limited cells captured in other studies as well as the use of different sequencing techniques |
|  | HLA-class II downregulation in CD14<sup>+</sup> monocytes (Fig. 5f) | Yes | Yes |  |
|  | Heterogenous ISG module upregulation in CD14<sup>+</sup> monocytes (Fig. 5g) | Yes | No | Patient-specific variability in fighting COVID-19 infection |
|  | Presence of type I IFN driven inflammatory signatures in CD14<sup>+</sup> monocytes (Fig. 5d, 5e) | Yes | Yes |  |
|  | Lack of substantial expression of pro-inflammatory cytokine genes (TNF, IL6, IL1B, CCL3, CCL4 or CXCL2) in CD14<sup>+</sup> monocytes (data not shown) | Yes | No | CD14<sup>+</sup> monocytes(FOSB) are not found in Wilk, but found in others (Zhang Lee, Chua). These cells express: TNF, IL1B, CCL3, CCL4, CXCL2 |
| Liao et al. <sup>9</sup> | Increased CD8<sup>+</sup> T-cell clonal expansion in moderate COVID-19 patients compared to severe (Fig. 6d; Supp Fig. S12b, S12c) | No | No | Patient-specific variability in fighting COVID-19 infection and lack of comparable stages across multiple datasets having TCR data |
| Chua et al. <sup>5</sup> | Activated macrophages expressing inflammatory chemokines including CCL2, CCL3, CCL20, CXCL1, CXCL3, CXCL10, IL8, IL1B and TNF in severe patients (data not shown) | Yes | No | Not CCL2, CCL3, CXCL10 are correct. Others are weak. Different studies captured different populations. |

## Materials and Methods

### Datasets

#### 10X Chromium healthy control

The 10X Chromium data for healthy PBMCs was downloaded from the 10X genomics website [https://support.10xgenomics.com/single-cell-gene-expression/datasets/3.1.0/5k_pbmc_NGSC3aggr]. This dataset includes samples from 8 Chromium Connect channels and 8 manual channels.

#### Wen dataset

The raw fastq files for the Wen dataset^4^ were downloaded from the GSA (Genome Sequence Archive) database^28,29^ under the accession number PRJCA002413, including all the files for scRNA-seq, TCR-seq and BCR-seq.

The fastq files were aligned to the human genome (GRCh38 version 3.0.0, including 33538 genes), which was downloaded from the 10X Chromium website (https://support.10xgenomics.com/single-cell-gene-expression/software/downloads/latest), using Cell Ranger (3.1.0). TCR and BCR results were aligned to the human vdj reference (version 4.0.0) from 10X Chromium website using Cell Ranger (3.1.0).

#### Liao dataset

For the Liao dataset^9^, the Cell Ranger mapped results together with the TCR results were downloaded from GEO^30^ under the accession number GSE145926. The downloaded scRNA-seq data includes 33538 features, which is the same as the reference genome used in the above mentioned datasets. The cell type annotation of the dataset was downloaded from GitHub [https://raw.githubusercontent.com/zhangzlab/covid_balf/master/all.cell.annotation.meta.txt]. The unannotated cells were considered as low quality and were excluded from analysis.

#### Lee dataset

Results for the Lee dataset^1^ was downloaded from GEO under the accession number GSE149689. The expression matrix also includes 33,538 features, which is the same as the reference genome used in the above-mentioned datasets.

#### Yu dataset

We downloaded the raw fastq files for the Yu dataset^7^ from the GSA (Genome Sequence Archive) database under the accession number PRJCA002579. Reads mapping was performed in the same way as on the Wen dataset.

#### Jiang dataset

The mapped results for the Jiang dataset^31^ was downloaded from figureshare [https://figshare.com/articles/dataset/Single_cell_and_immune_repertoire_profiling_of_COVID-19_patients_reveal_novel_therapeutic_candidates/12115095]. The expression matrix also includes 33,538 features, which is the same as the reference genome used in the above mentioned datasets.

#### Wilk dataset

We downloaded the raw fastq files of the Wilk dataset^2^ from the ENA database^32^ under the accession number PRJNA633393. According to Wilk et al., dropEst, samtools and STAR were used for the reads mapping. Human genome GRCh38 version 3.0.0 (the same as the Wen dataset reference) was used as the reference genome for reads mapping. The resulting expression matrix includes 33,538 features for 150,245 cells.

#### Zhang dataset

We downloaded the raw fastq files of the Zhang dataset^3^ from the GSA (Genome Sequence Archive) database under the accession number HRA000150. Reads mapping was performed in the same way as on the Wen dataset.

#### He dataset

We downloaded the mapped results of the He dataset^6^ from GEO under the accession number GSE147143. The expression matrix includes 33,578 features, 24,020 of which are overlapped with the features in the reference genome used in the Wen dataset.

#### Chua dataset

The mapped results of the Chua dataset^5^ were downloaded from figureshare [https://doi.org/10.6084/m9.figshare.12436517] as mentioned in the publication. The expression matrix includes 26,924 features, 18,410 of which are overlapped with the features in the reference genome used in the Wen dataset. The ‘celltype’ column in the dataset was used as cell type annotation.

### Data processing

#### Quality control

All the datasets include 33,538 features except the He and Chua datasets. For the He and Chua datasets, only the gene symbols that overlap with the other datasets are considered. All the datasets are combined together for quality control and downstream analyses. We filtered the cells with higher than 15% of mitochondrial contents. We also excluded cells of fewer than 500 UMIs or fewer than 200 features. Features expressed in fewer than 3 cells were removed in the analysis. We used Scrublet (version 0.2.1)^33^ to determine the doublets in the datasets. As Scrublet was designed to deal with 10X data, the result on Wilk dataset was not used. Cell clusters represented by doublets were removed from the analysis (**Supplementary Figure S14**). The cell numbers after quality control were listed in **Table 1**.

#### Data Integration

SCANPY^34^ workflow was used to analyze the data. The following steps were performed: data normalization, log-transformation, highly variable genes selection using the ‘cellranger’ flavor and principal component analysis. Then, we used Harmony (version 0.0.5)^8^ to integrate data from different samples. UMAP^35^ and Louvain^36^ clustering were calculated according to the Harmony corrected latent space.

#### Cell type annotation

For the eight datasets of the same reference genome (Wen, Liao, Lee, Yu and Jiang datasets) we combined the dataset to annotate all the cell types. We first used logistic regression in SCCAF^10^ to infer the cell types using the possible cell types. And each of the Louvain clusters was assigned to a cell type label accordingly. For Wilk dataset and Chua dataset, we used the cell type annotations adopted from the publication. For the He dataset, we annotated the cell types according to the marker gene expression.

To determine the cell types in the data, we first used a machine learning based method to generate cell type labels according to a reference dataset^2^, see **Methods**. First the platelets (PPBP, PF4) and epithelial cells (TPPP3, KRT18) clusters were excluded in the downstream analysis. According to the published marker genes, which have been summarized in **Supplementary Data 2**, we divided the data into three populations: Lymphoid cells (CD3D, CD3E, NKG7, NCAM1), Myeloid cells (CD68, CD14, FCGR3A, CD1C) and B cells (CD19, MS4A1, CD79A, MZB1).

#### Differential expression and Gene Ontology analysis

To account for the technical effects such as number of cells or sequencing depth, the hurdle model in MAST^37^ was used to model the differential expression of the cells. Only the genes with a false detection rate (FDR) lower than 0.01 was used for volcano plot and later gene ontology analysis.

In the MAST results, the genes with a log fold change value greater than 0 are considered as up-regulated genes, while the rest are the down-regulated genes. And the up-regulated genes in a cell cluster are used for gene ontology analysis. The python package of GProfiler^38^ was used to understand the pathway regulation. The gene module scores for HLA class II and ISG signature were calculated for each individual cell using sc.tl.score_genes function of scanpy^39^ v1.5.1.

#### TCR/BCR analysis

The TCR and BCR V(D)J data from the studies Wen et al. 2020^4^, Zhang et al. 2020^3^ and Liao et al. 2020^9^ (TCR only) were analyzed separately using the python library pyvdj^40^ v0.1.2. For TCR V(D)J data, only cells with at least one productive TRA and at least one productive TRB chain were considered for analysis. Similarly, for BCR V(D)J data, only cells with at least one productive IGH and at least one productive IGL or IGK chain were considered for analysis. The UMAP plots were generated using scanpy^39^ v1.5.1. The boxplots and barplots were generated using R-package ggplot2^41^ v3.3.2, ggpubr^42^ v0.4.0.999, rstatix^42,43^ v0.6.0.999 and tidyverse^44^ v1.3.0.

## Data Availability

For the other studies included in the current manuscript, the raw sequencing data are available in European Genome-phenome Archive (EGAS00001004481 for Chua et al.^5^), GEO (GSE149689 for Lee et al.^1^, GSE150728 for Wilk et al., GSE145926 for Liao et al.^9^) and Genome Sequence Archive (PRJCA002413 for Wen et al.^4^, PRJCA002564 for Zhang et al.^3^, PRJCA002579 for Yu et al.^7^, GSE147143 for He et al.^6^). Details are listed in **Table 1**.

The merged dataset can be visualised interactively through cellxgene in the Human Cell Atlas Galaxy.EU instance^45^, following instructions in the Supplementary Material Note 1.

## Acknowledgements

We thank Prof. Catherine Blish for providing us with WHO scores^46^ of samples from Wilk et al. 2020^2^ that helped us to redefine non-ventilated patients as moderate for comparison. We would like to acknowledge Dr. Pedro Beltrao and other members of the EMBL COVID-19 discussion group for useful suggestions. We also thank Dr. Craig Russell for comments.

## Author contributions

MG analysed the data and wrote the paper. XL summarized patient level clinical data, gave biological inputs and wrote the paper. ZM designed the project, collected the datasets, analysed the data and wrote the paper. PM and IP visualised the single cell data. AB, ZM and YS supervised the project.

## References

1. Lee, J. S. et al. Immunophenotyping of COVID-19 and influenza highlights the role of type I interferons in development of severe COVID-19. Sci Immunol 5, (2020).

2. Wilk, A. J. et al. A single-cell atlas of the peripheral immune response in patients with severe COVID-19. Nat. Med. 26, 1070–1076 (2020).

3. Zhang, J.-Y. et al. Single-cell landscape of immunological responses in patients with COVID-19. Nat. Immunol. 21, 1107–1118 (2020).

4. Wen, W. et al. Immune cell profiling of COVID-19 patients in the recovery stage by single-cell sequencing. Cell Discov 6, 31 (2020).

5. Chua, R. L. et al. COVID-19 severity correlates with airway epithelium-immune cell interactions identified by single-cell analysis. Nat. Biotechnol. 38, 970–979 (2020).

6. He, J. et al. Single-cell analysis reveals bronchoalveolar epithelial dysfunction in COVID-19 patients. Protein Cell 11, 680–687 (2020).

7. Yu, K. et al. Thymosin alpha-1 Protected T Cells from Excessive Activation in Severe COVID-19. (2020) doi:10.21203/rs.3.rs-25869/v1.

8. Korsunsky, I. et al. Fast, sensitive and accurate integration of single-cell data with Harmony. Nat. Methods 16, 1289–1296 (2019).

9. Liao, M. et al. Single-cell landscape of bronchoalveolar immune cells in patients with COVID-19. Nature Medicine vol. 26 842–844 (2020).

10. Miao, Z. et al. Putative cell type discovery from single-cell gene expression data. Nat. Methods 17, 621–628 (2020).

11. Li, H. et al. Dysfunctional CD8 T Cells Form a Proliferative, Dynamically Regulated Compartment within Human Melanoma. Cell vol. 181 747 (2020).

12. Künkele, K.-P. et al. Vγ9Vδ2 T Cells: Can We Re-Purpose a Potent Anti-Infection Mechanism for Cancer Therapy? Cells 9, (2020).

13. Oliver Nussbaumer, M. T. Functional Phenotypes of Human Vγ9Vδ2 T Cells in Lymphoid Stress Surveillance. Cells 9, (2020).

14. Schirmer, L., Rothhammer, V., Hemmer, B. & Korn, T. Enriched CD161high CCR6+ yδT cells in the cerebrospinal fluid of patients with multiple sclerosis. JAMA Neurol. 70, 345–351 (2013).

15. Glatzel, A. et al. Patterns of chemokine receptor expression on peripheral blood gammadelta T lymphocytes: strong expression of CCR5 is a selective feature of V delta 2/Vgamma 9 gamma delta T cells. J. Immunol. 168, (2002).

16. Park, D. et al. Differences in the molecular signatures of mucosal-associated invariant Tcells and conventional T cells. Sci. Rep. 9, (2019).

17. Wong, E. B. et al. TRAV1-2 + CD8 + T-cells including oligoconal expansions of MAITcells are enriched in the airways in human tuberculosis. Communications Biology 2, 1–13 (2019).

18. Schulte-Schrepping, J. et al. Severe COVID-19 Is Marked by a Dysregulated Myeloid Cell Compartment. Cell 182, 1419–1440.e23 (2020).

19. Wong, W. S. et al. Reference ranges for lymphocyte subsets among healthy Hong Kong Chinese adults by single-platform flow cytometry. Clin. Vaccine Immunol. 20, 602–606 (2013).

20. Zhang, F. et al. Adaptive immune responses to SARS-CoV-2 infection in severe versus mild individuals. Signal Transduction and Targeted Therapy vol. 5 (2020).

21. [No title]. https://www.who.int/blueprint/priority-diseases/key-action/COVID-19_Treatment_Trial_Design_Master_Protocol_synopsis_Final_18022020.pdf?ua=1.

22. Lin, S.-J., Kuo, M.-L., Hsiao, H.-S. & Lee, P.-T. Azithromycin modulates immune response of human monocyte-derived dendritic cells and CD4+ T cells. Int.Immunopharmacol. 40, 318–326 (2016).

23. Su, Y. et al. Multi-Omics Resolves a Sharp Disease-State Shift between Mild and Moderate COVID-19. Cell 183, 1479–1495.e20 (2020).

24. Overmyer, K. A. et al. Large-Scale Multi-omic Analysis of COVID-19 Severity. Cell Syst 12, 23–40.e7 (2021).

25. Bernardes, J. P. et al. Longitudinal Multi-omics Analyses Identify Responses of Megakaryocytes, Erythroid Cells, and Plasmablasts as Hallmarks of Severe COVID-19. Immunity 53, 1296–1314.e9 (2020).

26. Barh, D. et al. Multi-omics-based identification of SARS-CoV-2 infection biology and candidate drugs against COVID-19. Comput. Biol. Med. 126, 104051 (2020).

27. Stephenson, E. et al. The cellular immune response to COVID-19 deciphered by single cell multi-omics across three UK centres. bioRxiv (2021)doi:10.1101/2021.01.13.21249725.

28. Wang, Y. et al. GSA: Genome Sequence Archive<sup>. Genomics Proteomics Bioinformatics 15, 14–18 (2017).

29. National Genomics Data Center Members and Partners. Database Resources of theNational Genomics Data Center in 2020. Nucleic Acids Res. 48, D24–D33 (2020).

30. Barrett, T. et al. NCBI GEO: archive for functional genomics data sets--update. NucleicAcids Res. 41, D991–5 (2013).

31. Jiang, Q. Single cell and immune repertoire profiling of COVID-19 patients reveal novel therapeutic candidates. (2020) doi:10.5281/zenodo.3747336.

32. Amid, C. et al. The European Nucleotide Archive in 2019. Nucleic Acids Res. 48, D70–D76 (2020).

33. Wolock, S. L., Lopez, R. & Klein, A. M. Scrublet: Computational Identification of Cell Doublets in Single-Cell Transcriptomic Data. Cell Syst 8, 281–291.e9 (2019).

34. Wolf, F. A., Angerer, P. & Theis, F. J. SCANPY: large-scale single-cell gene expression data analysis. Genome Biol. 19, 15 (2018).

35. Becht, E. et al. Dimensionality reduction for visualizing single-cell data using UMAP. Nat. Biotechnol. 37, 38–44 (2018).

36. Blondel, V. D., Guillaume, J.-L., Lambiotte, R. & Lefebvre, E. Fast unfolding of communities in large networks. Journal of Statistical Mechanics: Theory and Experiment vol. 2008 P10008 (2008).

37. Finak, G. et al. MAST: a flexible statistical framework for assessing transcriptional*changes and characterizing heterogeneity in single-cell RNA sequencing data*. Genome Biology vol. 16 (2015).

38. Raudvere, U. et al. g:Profiler: a web server for functional enrichment analysis and conversions of gene lists (2019 update). Nucleic Acids Res. 47, W191–W198 (2019).

39. Wolf, F. A., Angerer, P. & Theis, F. J. SCANPY: large-scale single-cell gene expression data analysis. Genome Biol. 19, 15 (2018).

40. veghp. veghp/pyVDJ. https://github.com/veghp/pyVDJ.

41. Wickham, H. ggplot2: Elegant Graphics for Data Analysis. (Springer Science & Business Media, 2009).

42. Website. kassambara. kassambara/ggpubr. https://github.com/kassambara/ggpubr.

43. Website. kassambara. kassambara/rstatix. https://github.com/kassambara/rstatix.

44. Wickham, H. et al. Welcome to the Tidyverse. Journal of Open Source Software 4, 1686 (2019).

45. Moreno, P. et al. User-friendly, scalable tools and workflows for single-cell analysis. Cold Spring Harbor Laboratory 2020.04.08.032698 (2020) doi:10.1101/2020.04.08.032698.

46. [No title]. https://www.who.int/blueprint/priority-diseases/key-action/COVID-19_Treatment_Trial_Design_Master_Protocol_synopsis_Final_18022020.pdf?ua=1.

