## Supplementary File for "Meta-analysis reveals consistent immune response patterns in COVID-19 infected patients at single-cell resolution"

### Supplementary Tables

#### Supplementary Table S1. Simplification of the stage definition

| Dataset | Stage annotation in original study | Stage annotation in current study | Any additional comment |
|---|---|---|---|
| 10X | Healthy | Healthy | |
| Lee et al. | Asymptomatic | Asymptomatic | |
| | Healthy | Healthy | |
| | Influenza | Influenza | |
| | Mild | Mild | This is kept as Mild, as it shows a different distribution from other moderate samples. |
| | Severe | Severe | |
| Wilk et al. | Healthy | Healthy | |
| | NonVent | Moderate | |
| | Vent | Severe | |
| Zhang et al. | Healthy | Healthy | |
| | Moderate | Moderate | |
| | Severe | Severe | |
| | convalescent | convalescent | |
| Wen et al. | Healthy | Healthy | |
| | early recovery | convalescent | |
| | late recovery | late recovery | |
| Yu et al. | Healthy | Healthy | |
| | Convalescence Mild | convalescent | |
| | P&C Mild | Post Mild | |
| | Post Mild | Post Mild | |
| Jiang et al. | early recovery | convalescent | |
| Liao et al. | Healthy | Healthy | |
| | Moderate | Moderate | |
| | Severe | Severe | |
| He et al. | Severe | Severe | |
| Chua et al. | Healthy | Healthy | |
| | Moderate | Moderate | |
| | critical | Severe | |

Supplementary Table S2. The self-projection accuracy of the 5 cell populations.

| Self-projection accuracy | Cross_validation | Training set | Test set |
|---|---|---|---|
| **B** | 0.9535 | 0.9608 | 0.9200 |
| **Lymphoid** | 0.9509 | 0.9671 | 0.9492 |
| **Myeloid** | 0.9567 | 0.9688 | 0.9404 |
| **Epithelial** | 0.9873 | 0.9922 | 0.9863 |
| **Platelets** | 0.9934 | 0.9970 | 0.9957 |

### Supplementary Figures

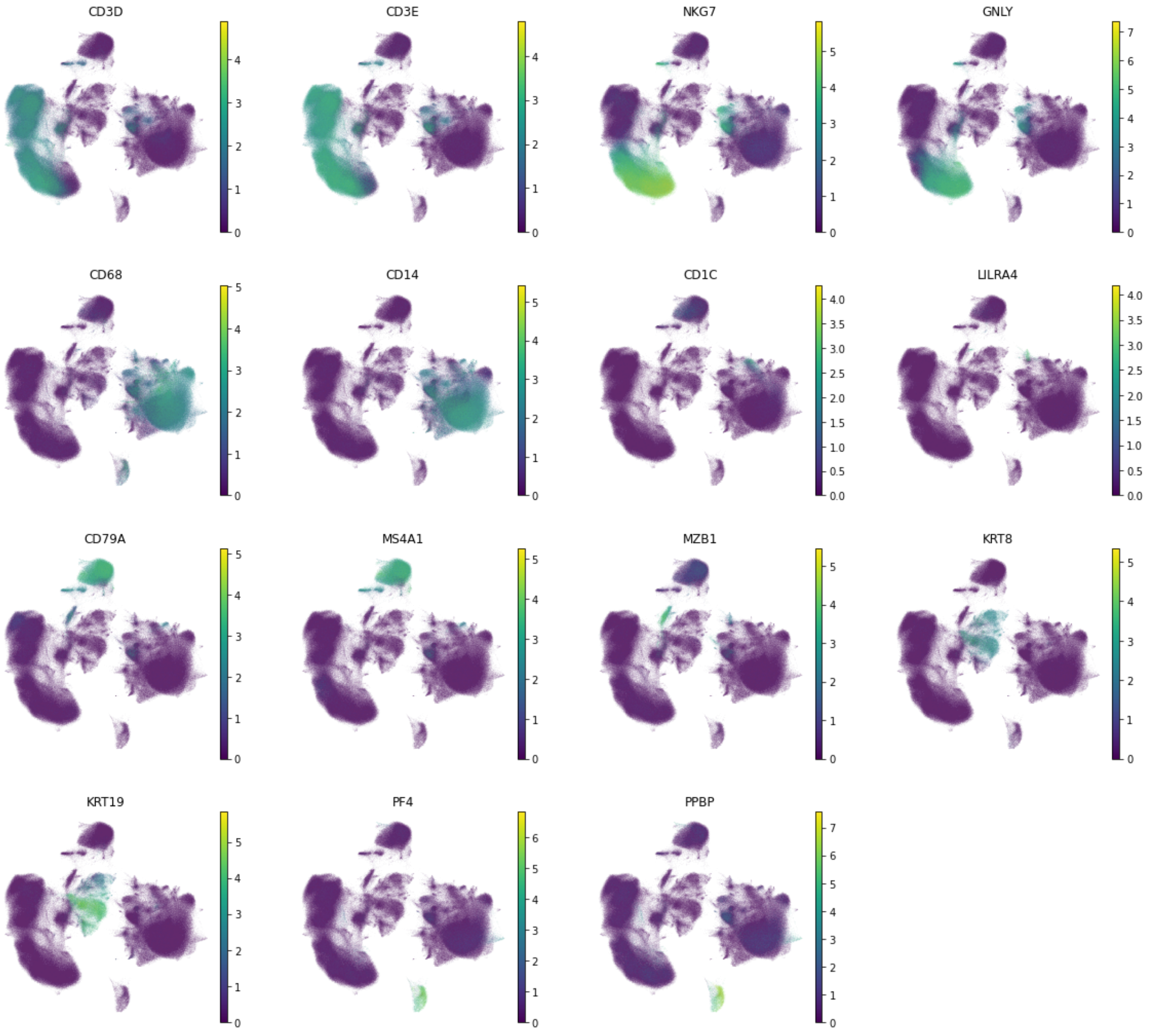


#### Supplementary Figure S1. Marker gene expression for the 5 cell populations.

The UMAPs show the marker gene expression of Lymphoid cells (CD3D, CD3E, NKG7, GNLY), Myeloid cells(CD68, CD14, CD1C, LILRA4), B cells (CD79A, MS4A1, MZB1), Epithelial cells (KRT8, KRT19) and Platelets (PF4, PPBP).

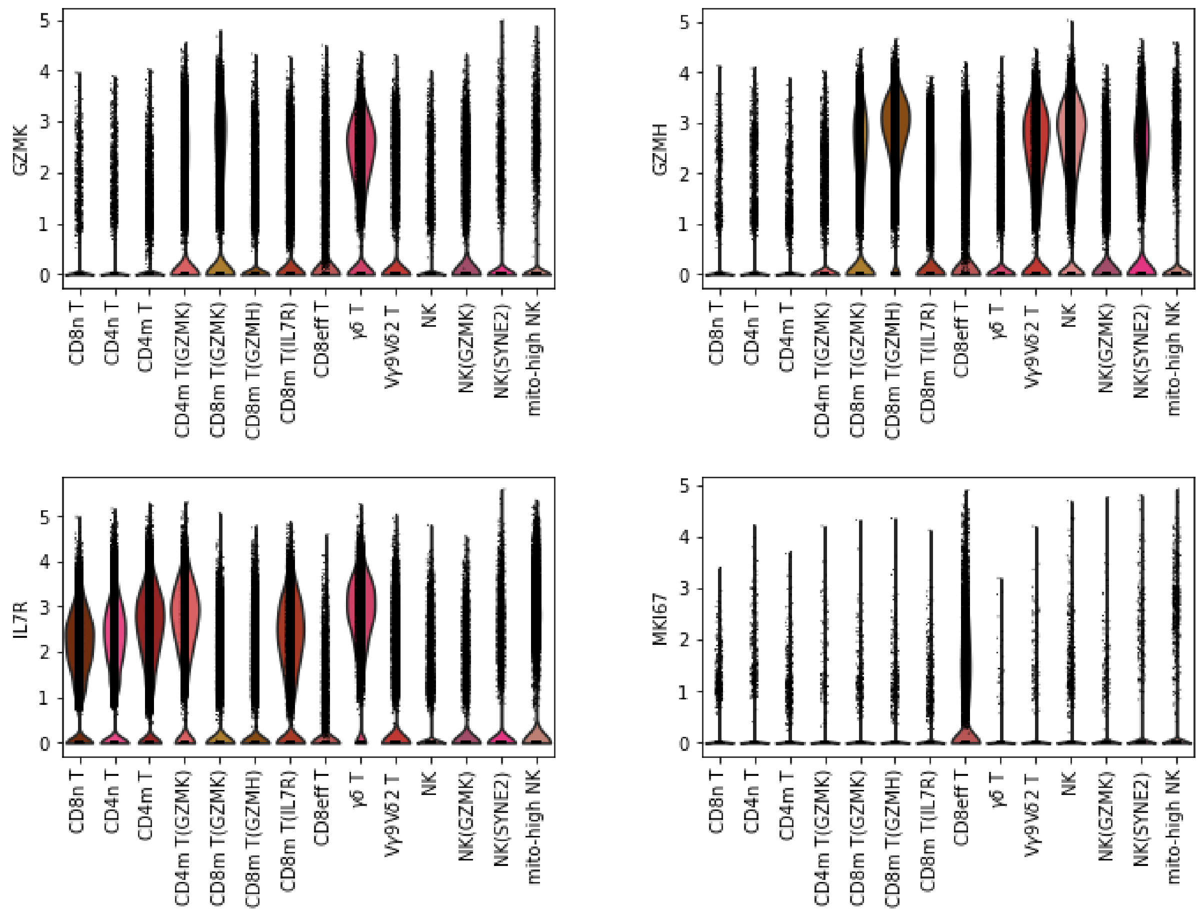


Supplementary Figure S2. The violin plots of the marker gene expressions for CD8 T cell subpopulations.

GZMK, GZMH, IL7R and MKI67 are used to distinguish CD8m T(GZMK), CD8m T(GZMH), CD8m T(IL7R) and CD8eff T cells.

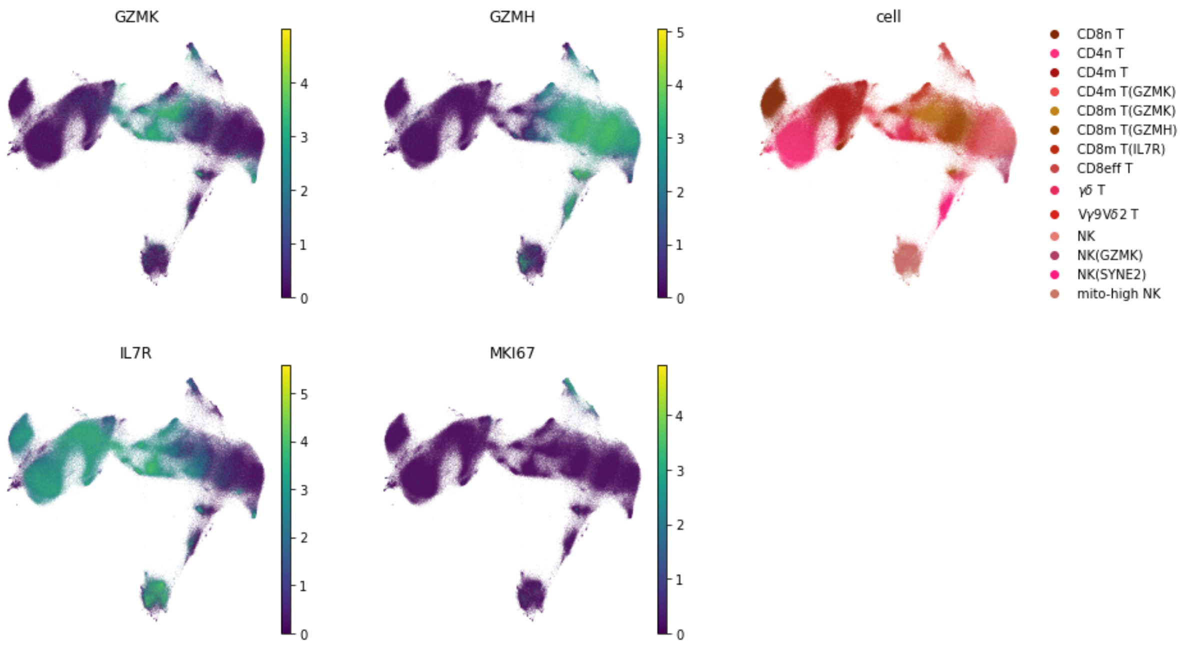


#### Supplementary Figure S3. The UMAP plots of the marker gene expressions for CD8 T cell populations

GZMK, GZMH, IL7R and MKI67 are used to distinguish CD8m T(GZMK), CD8m T(GZMH), CD8m T(IL7R) and CD8eff T cells. GZMH and GZMK not only show good discrimination of the CD8 T cell subpopulations, but also discriminates the NK cell populations.

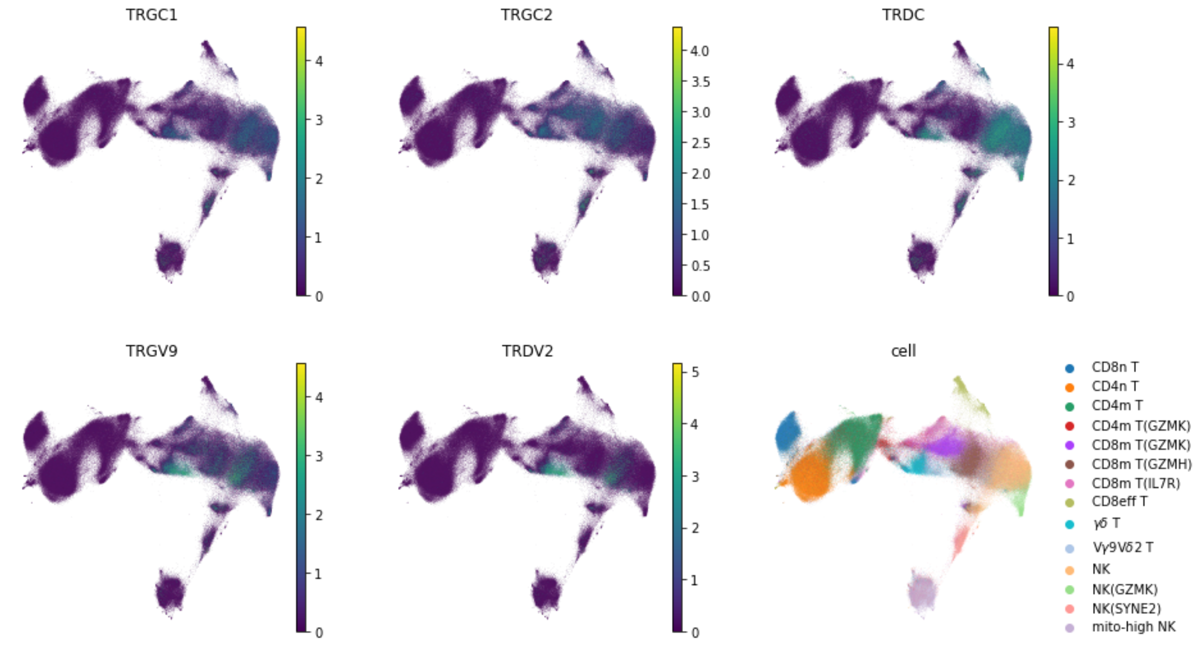


Supplementary Figure S4. The UMAP plots of the marker gene expressions for γδ T cell populations

TRGC1, TRGC2 and TRDC are the marker genes suggested by Wilk et al. , while TRGV9 and TRDV2 are suggested by Zhang et al.

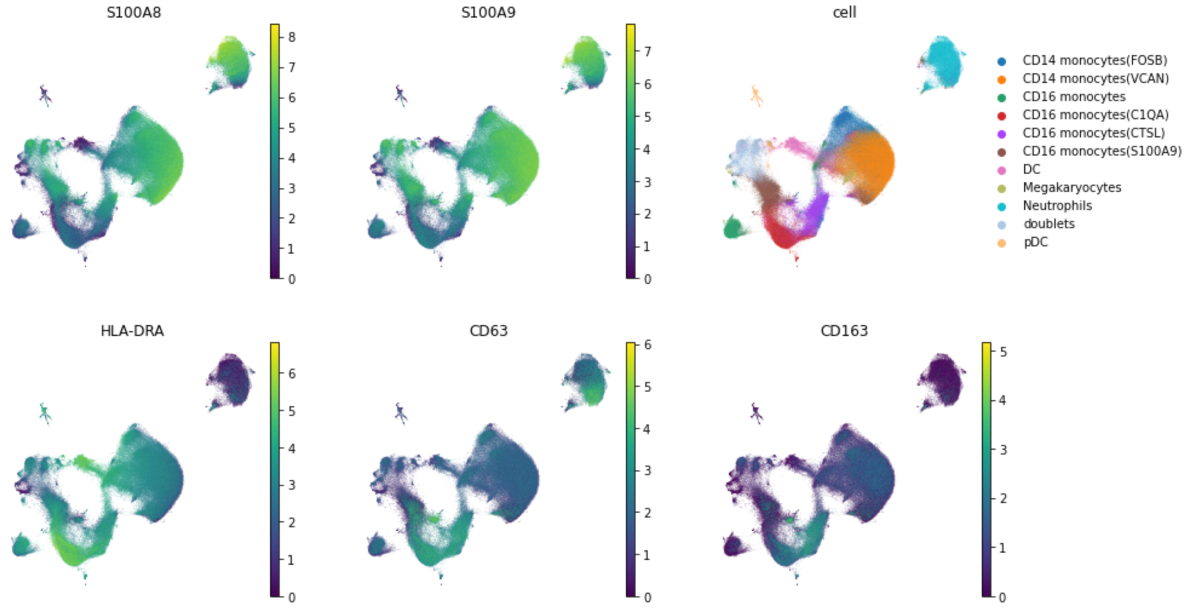


Supplementary Figure S5. The UMAP plots of the marker gene expressions for CD16 monocytes populations.

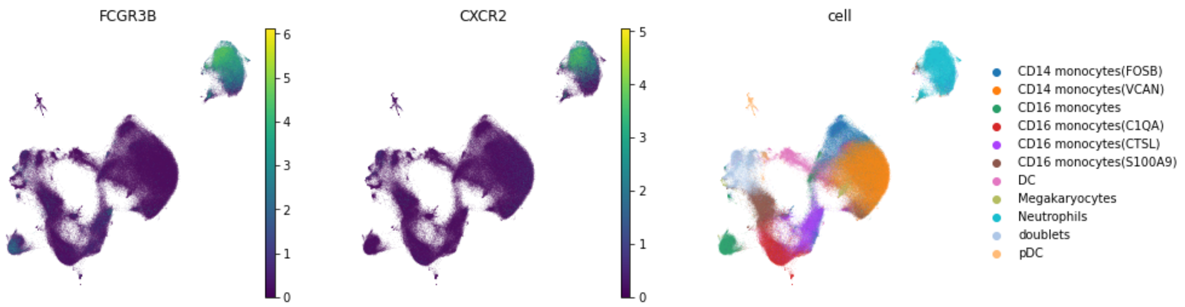


Supplementary Figure S6. The UMAP plots of the marker gene expressions for neutrophils.

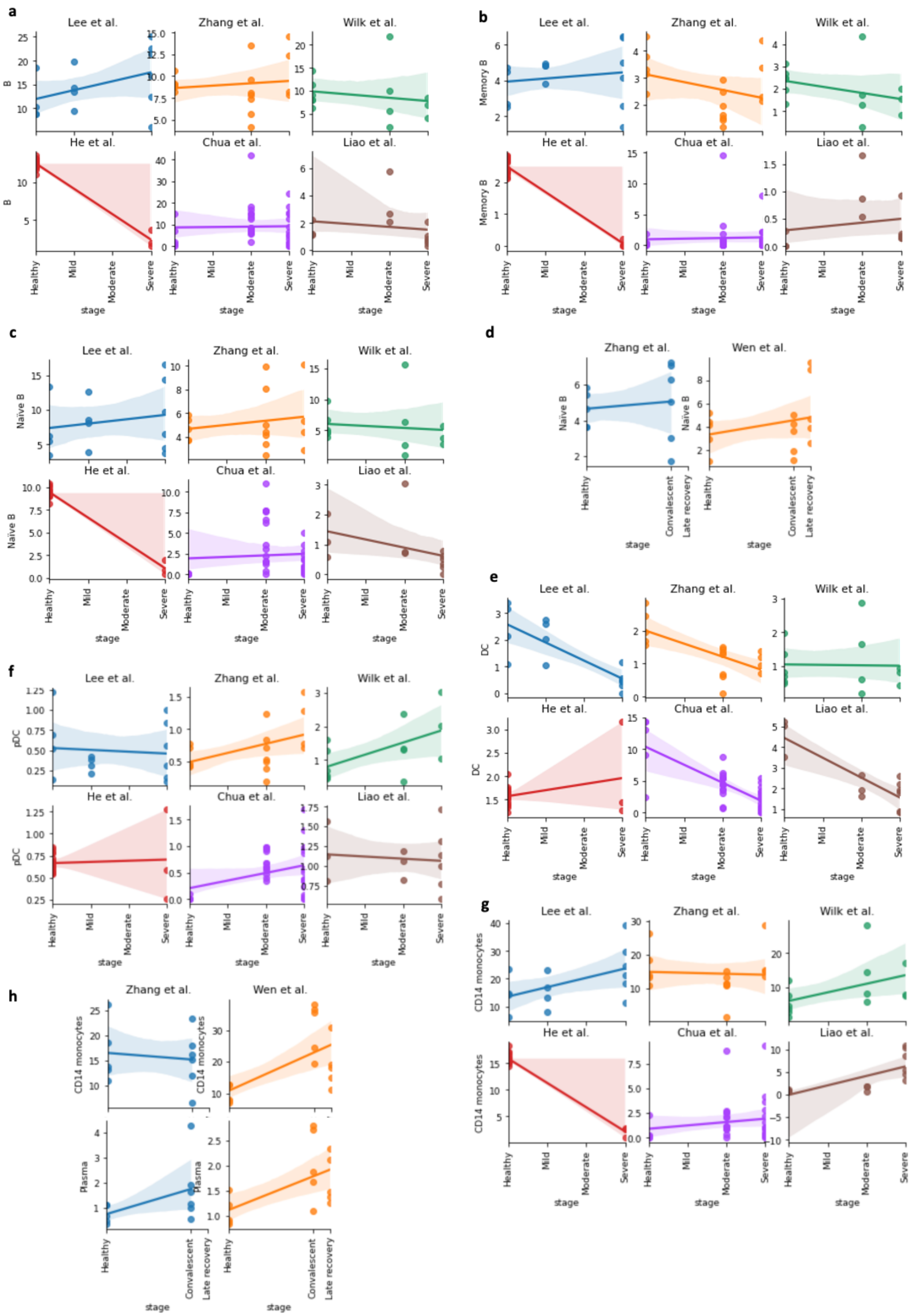


Supplementary Figure S7. The cell proportion changes in a) all B cells; b) memory B-cells; c) naïve B-cells across datasets with stages healthy, (mild), moderate and severe; d) naïve B-cells across datasets with stages healthy, convalescent (and late recovery); e) dendritic cells (DC); f) Plasmacytoid dendritic cells (pDC); g) CD14$^{+}$ monocytes; h) plasma cells and CD14$^{+}$ monocytes with stages healthy, convalescent (and late recovery)

CD4 T

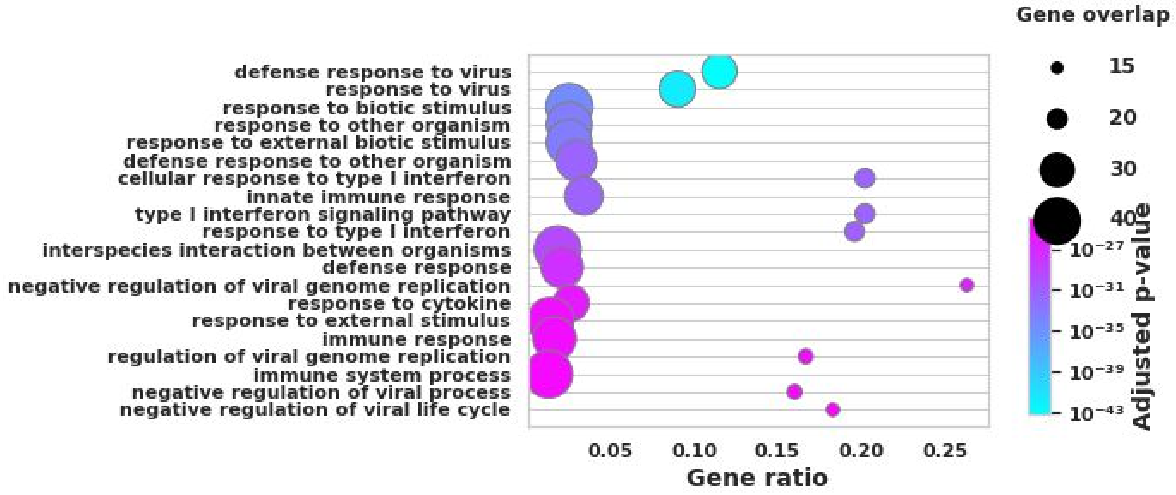


CD8 T

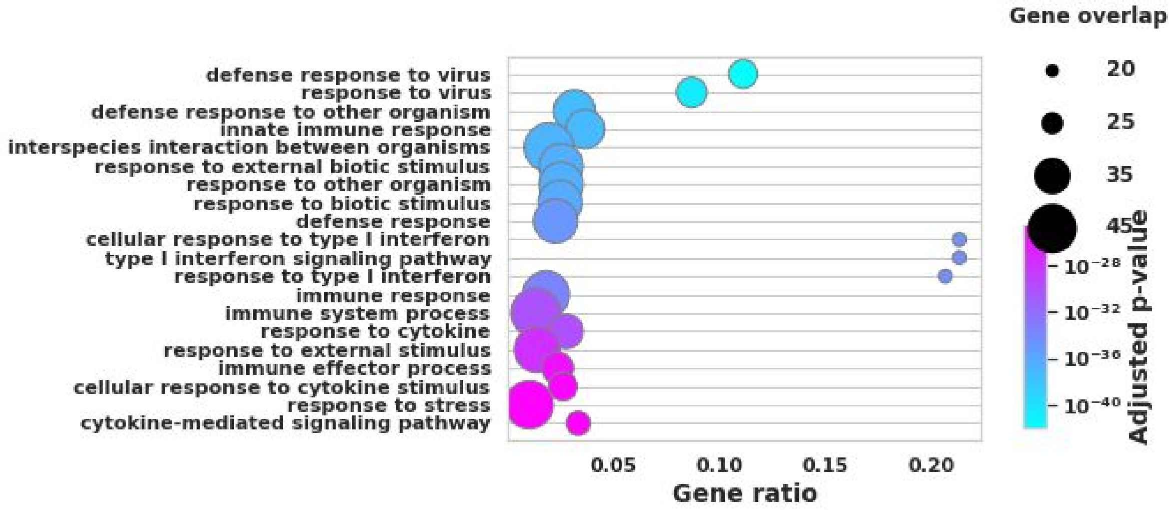


γδ T

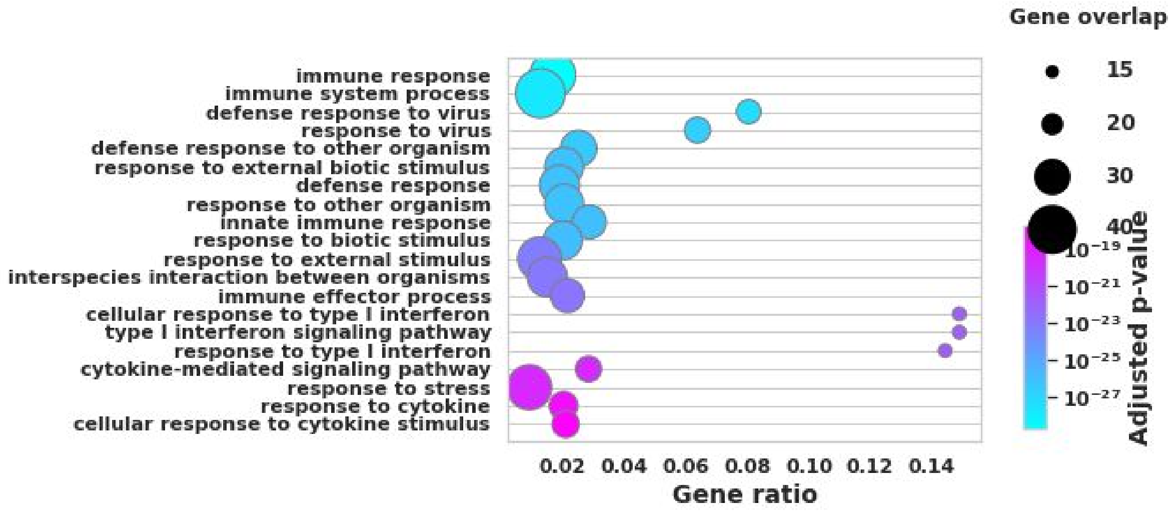


DC

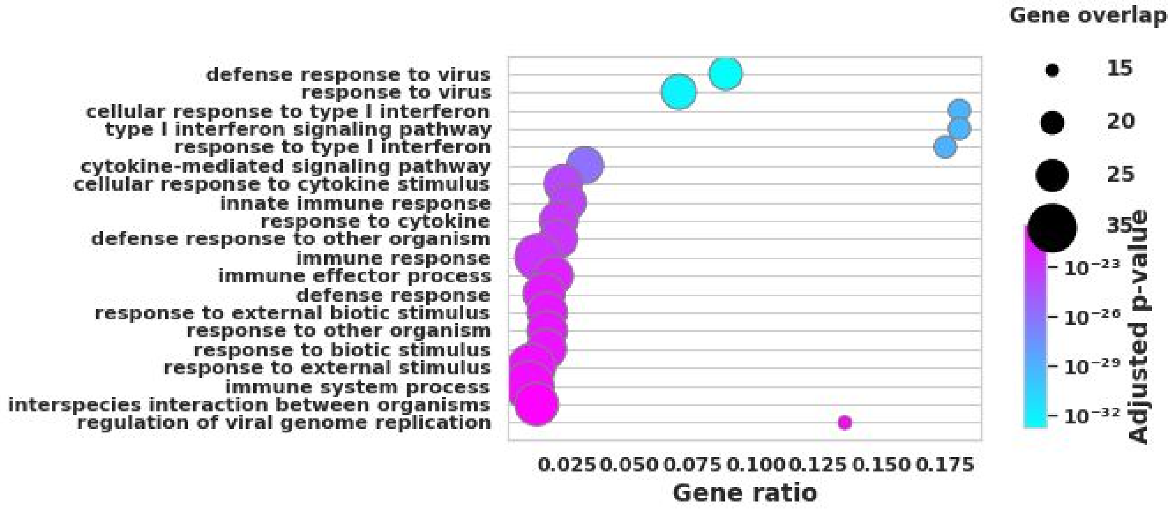


pDC

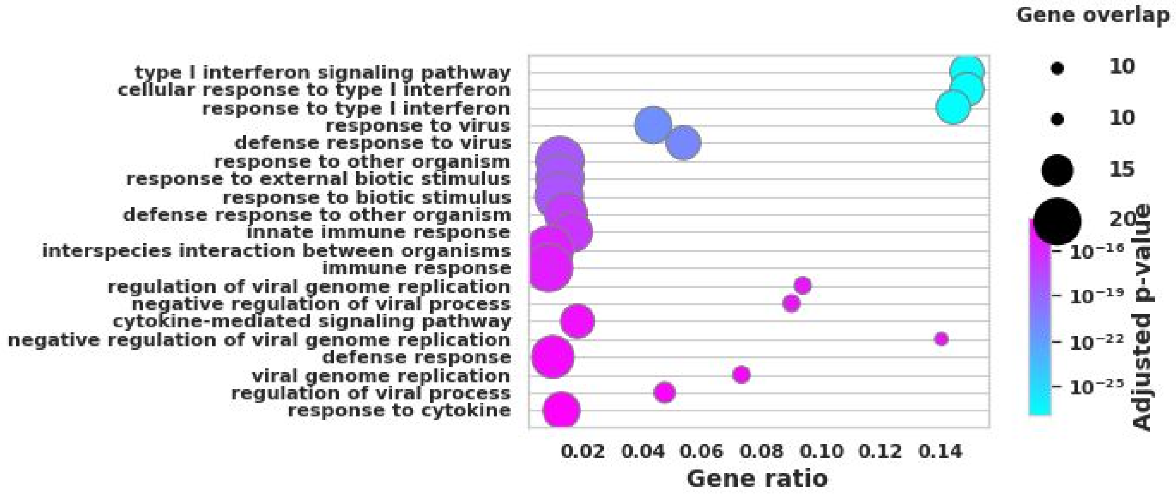


Neutrophils

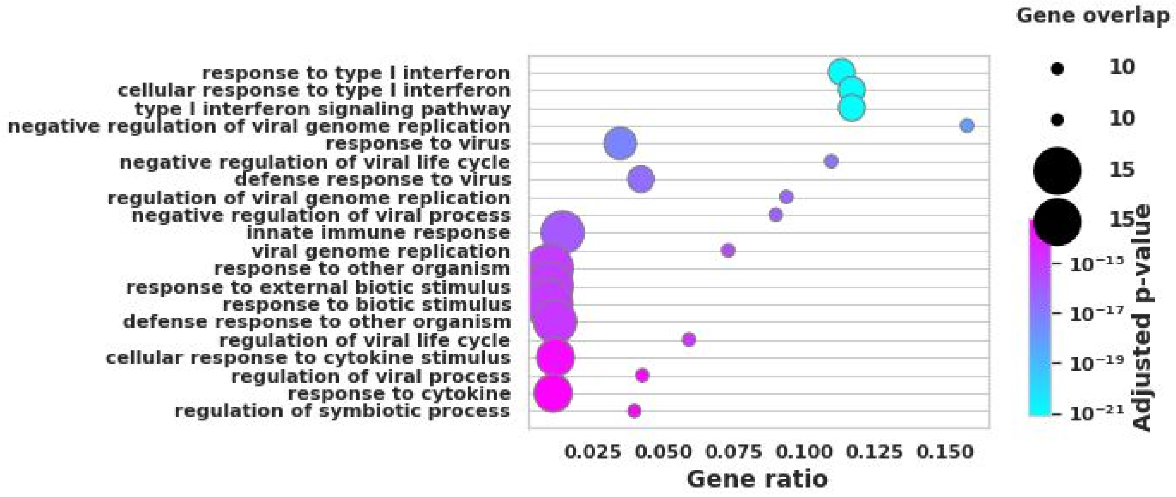
Gene overlap
response to type I interferon
cellular response to type I interferon
type I interferon signaling pathway
negative regulation of viral genome replication
response to virus
negative regulation of viral life cycle
defense response to virus
regulation of viral genome replication
negative regulation of viral process
innate immune response
viral genome replication
response to other organism
response to external biotic stimulus
response to biotic stimulus
defense response to other organism
regulation of viral life cycle
cellular response to cytokine stimulus
regulation of viral process
response to cytokine
regulation of symbiotic process
0.025 0.050 0.075 0.100 0.125 0.150
Gene ratio
10
10
15
15
$10^{-15}$
$10^{-17}$
$10^{-19}$
$10^{-21}$
Adjusted p-value

Plasma

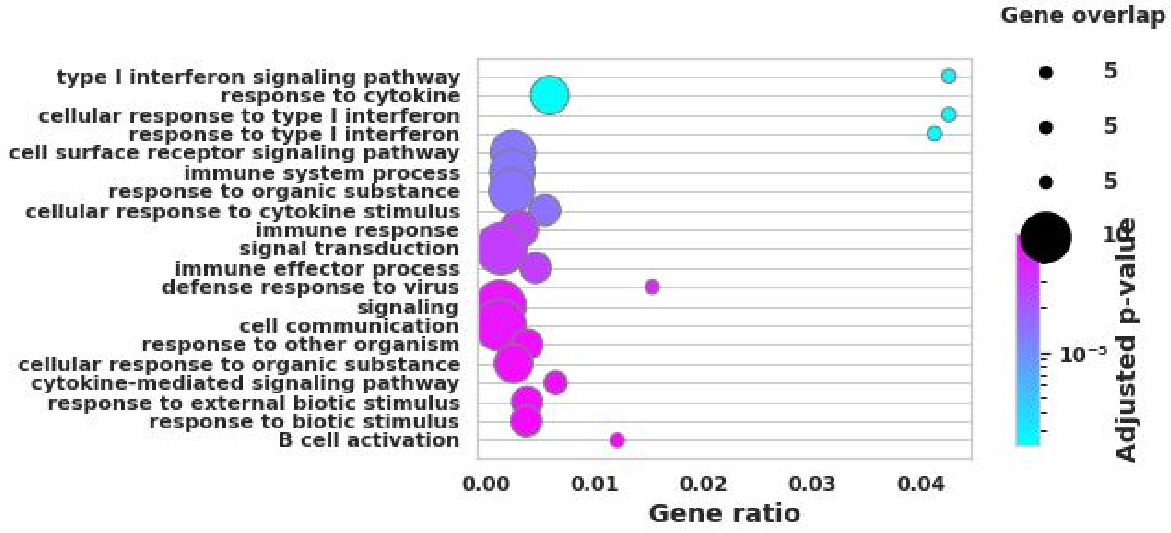
Gene overlap
type I interferon signaling pathway
response to cytokine
cellular response to type I interferon
response to type I interferon
cell surface receptor signaling pathway
immune system process
response to organic substance
cellular response to cytokine stimulus
immune response
signal transduction
immune effector process
defense response to virus
signaling
cell communication
response to other organism
cellular response to organic substance
cytokine-mediated signaling pathway
response to external biotic stimulus
response to biotic stimulus
B cell activation
0.00 0.01 0.02 0.03 0.04
Gene ratio
5
5
5
$10^{-5}$
Adjusted p-value

B

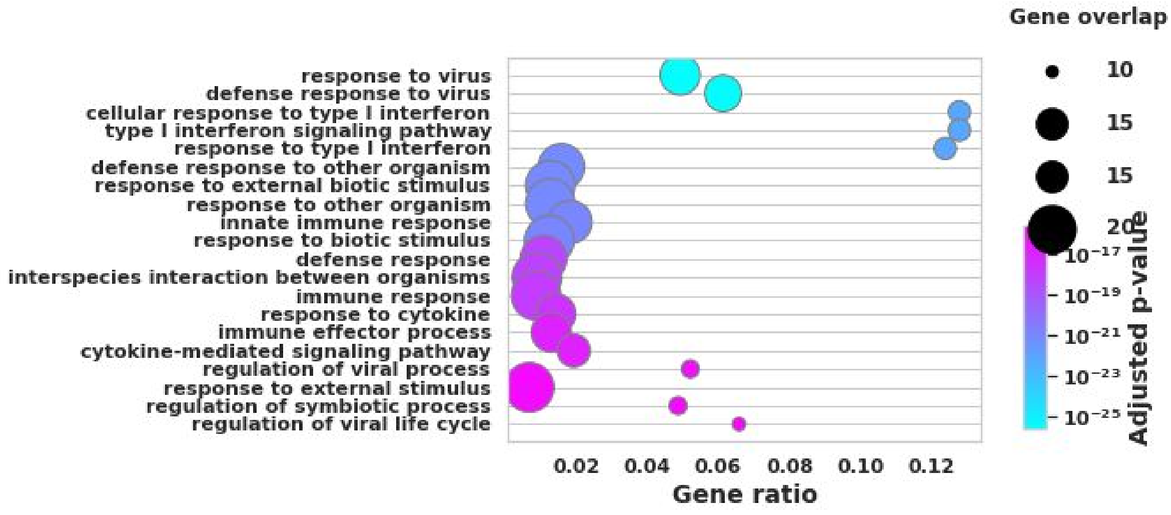
Gene overlap
response to virus
defense response to virus
cellular response to type I interferon
type I interferon signaling pathway
response to type I interferon
defense response to other organism
response to external biotic stimulus
response to other organism
innate immune response
response to biotic stimulus
defense response
interspecies interaction between organisms
immune response
response to cytokine
immune effector process
cytokine-mediated signaling pathway
regulation of viral process
response to external stimulus
regulation of symbiotic process
regulation of viral life cycle
0.02 0.04 0.06 0.08 0.10 0.12
Gene ratio
10
15
15
20
$10^{-17}$
$10^{-19}$
$10^{-21}$
$10^{-23}$
$10^{-25}$
Adjusted p-value

NK

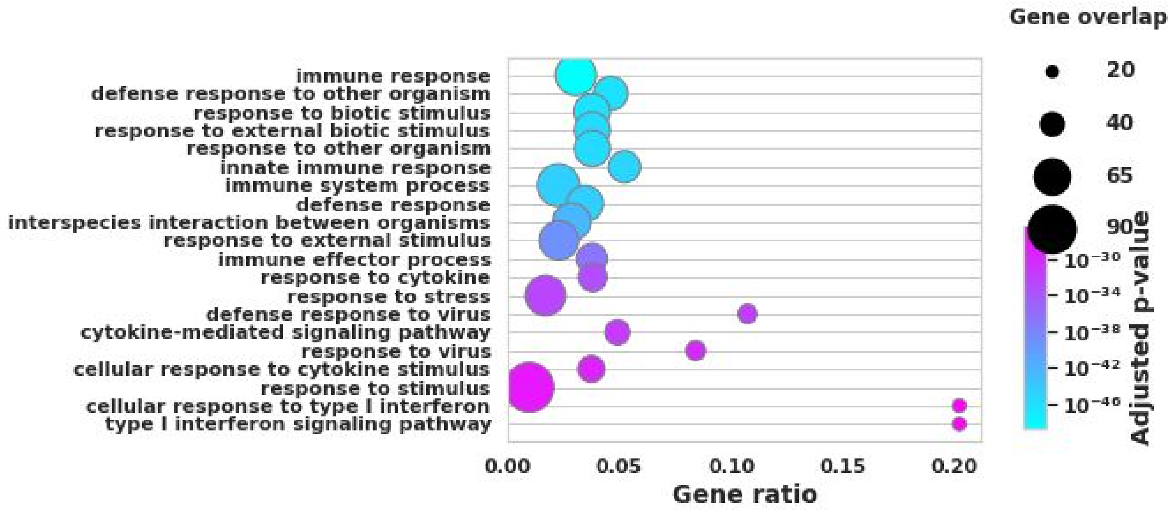


Supplementary Figure S8. Dot plots showing the top upregulated pathways traced from the overlapping differentially expressed (DE) genes (false discovery rate < 0.01) across multiple datasets in each cell-type.

These DE genes were found to be upregulated in both moderate vs healthy control comparison and severe vs healthy control comparison. They were further identified to be in common when the same analysis was performed in multiple datasets (Zhang, Wilk and Liao datasets)

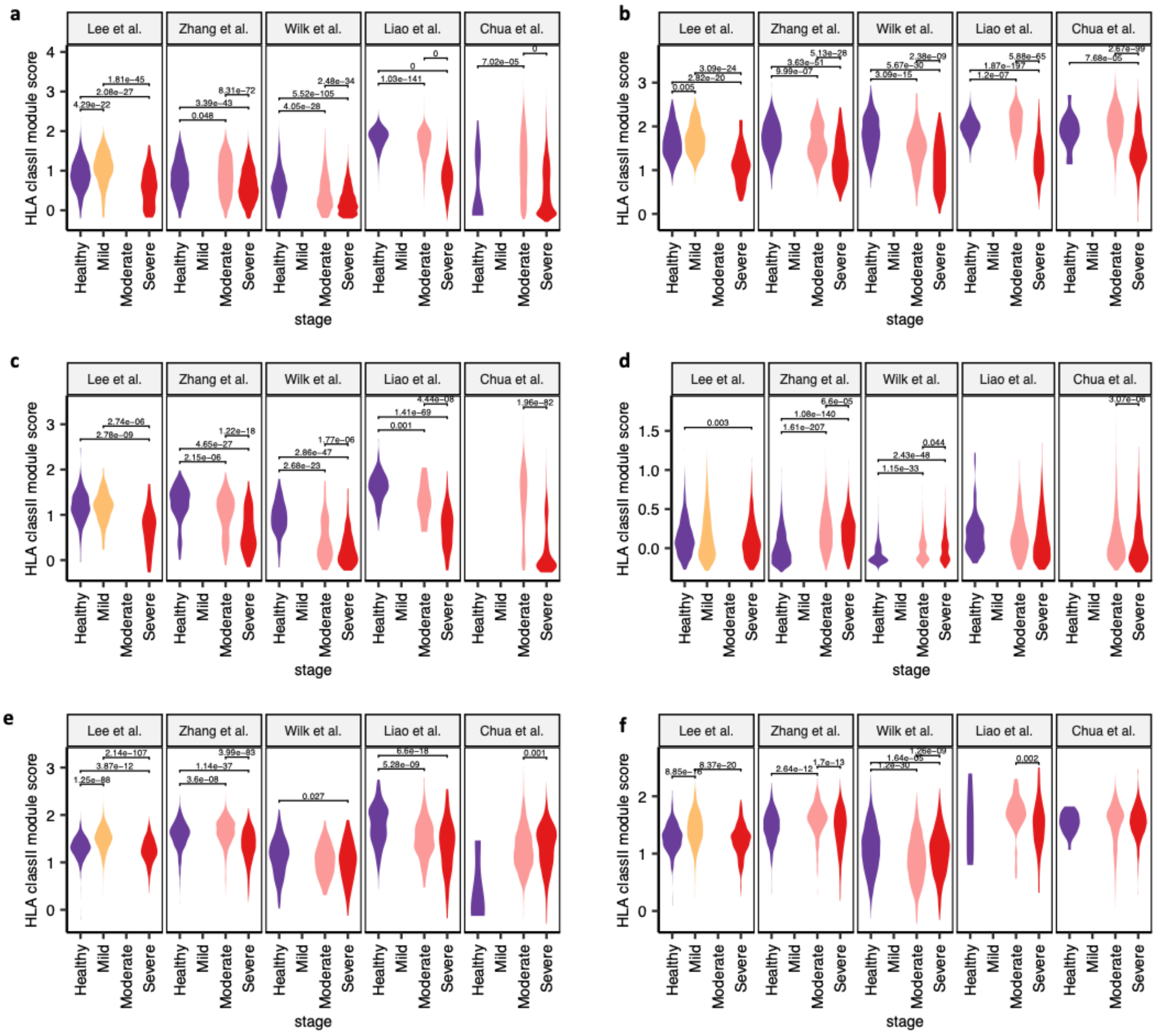


Supplementary Figure S9. Significant HLA class II downregulation in severe COVID-19 patients compared to healthy controls was observed in multiple cell-types across studies

- **a)** CD16+ monocytes
- **b)** Dendritic cells
- **c)** Plasmacytoid dendritic cells
- **d)** Natural killer cells
- **e)** Naive B-cells
- **f)** Memory B-cells

All differences were analyzed using two-sided unpaired Wilcoxon rank sum tests with Bonferroni correction and p-values <0.05 are reported.

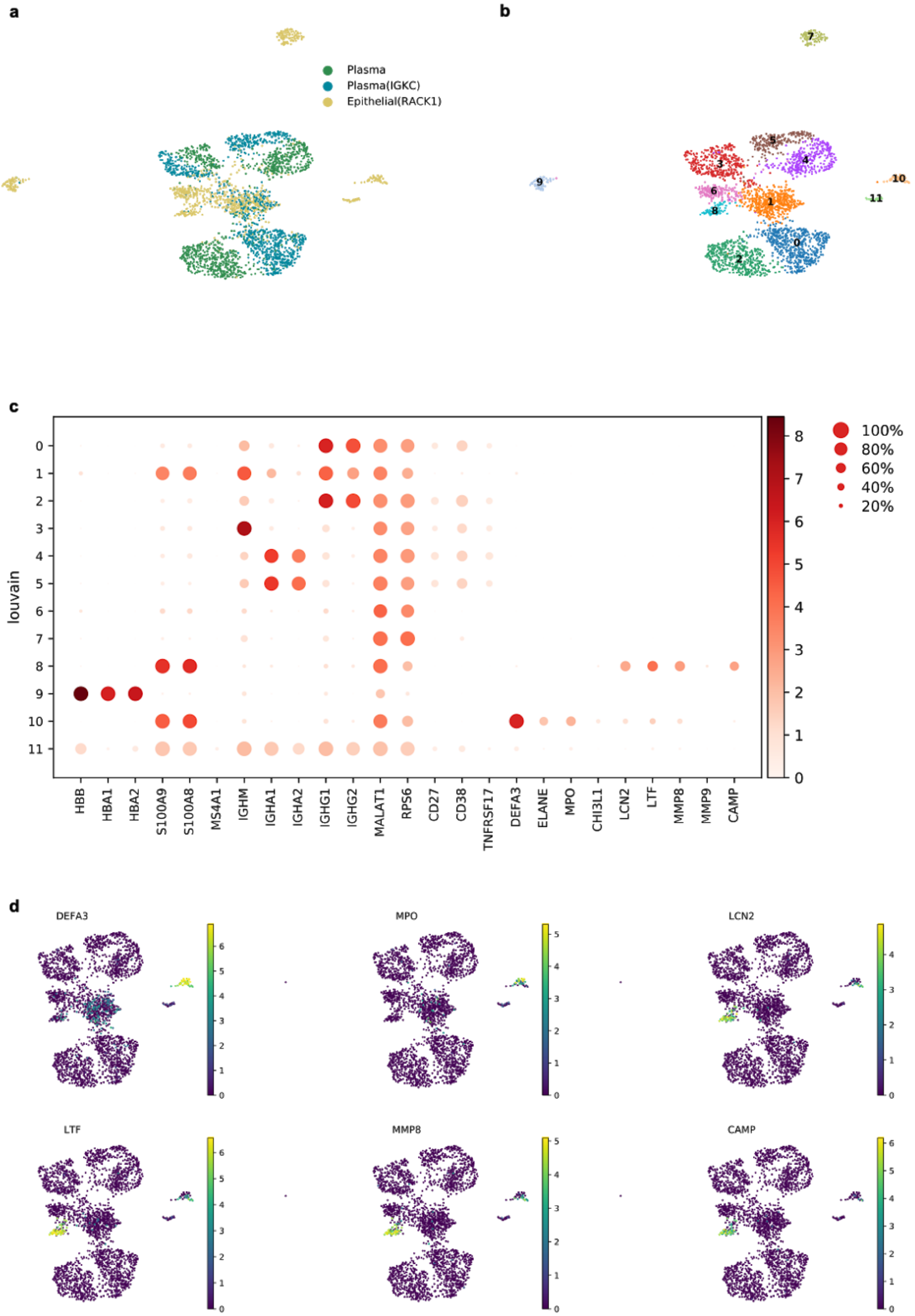
a
b
Plasma
Plasma(IGKC)
Epithelial(RACK1)
c
louvain
HBB
HBA1
HBA2
S100A9
S100A8
MS4A1
IGHM
IGHA1
IGHA2
IGHG1
IGHG2
MALAT1
RPS6
CD27
CD38
TNFRSF17
DEFA3
ELANE
MPO
CHI3L1
LCN2
LTF
MMP8
MMP9
CAMP
100%
80%
60%
40%
20%
d
DEFA3
MPO
LCN2
LTF
MMP8
CAMP

Supplementary Figure S10. Reproduction of developing neutrophil population in severe COVID-19 patients from Wilk et al. 2020 dataset.

a) UMAP embedding showing the plasma cell and epithelial cell (RACK1) subpopulations in the severe COVID-19 patients of Wilk et al. 2020 dataset (n=2862 cells)
b) UMAP embedding showing the results of louvain clustering on panel a)
c) Dotplot showing the marker gene expression in each of the corresponding louvain clusters shown on the y-axis.
d) UMAP embedding colored by selected developing neutrophil marker genes' expression where primary granule proteins are encoded by *DEFA3* and *MPO*, secondary granule proteins are encoded by *LCN2* and *LTF* and the tertiary granule proteins are encoded by *MMP8* and *CAMP* as used by Wilk et al. 2020. Note that the louvain cluster 7 and 9 have been removed from this representation to improve the graphics as they don't represent either plasma cells or developing neutrophil population (panel c).

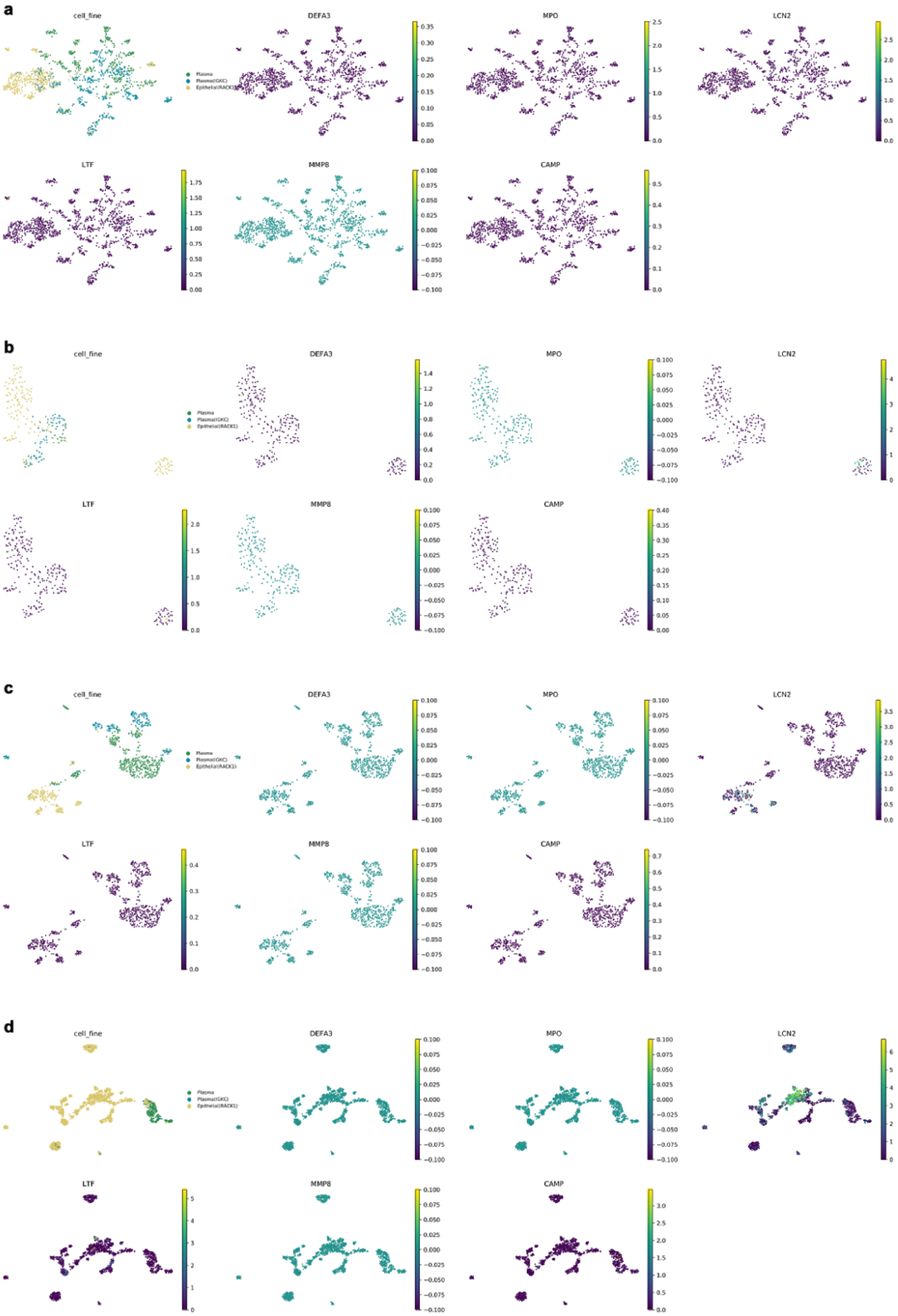
a
cell_fine
DEFA3
MPO
LCN2
LTF
MMP8
CAMP
b
cell_fine
DEFA3
MPO
LCN2
LTF
MMP8
CAMP
c
cell_fine
DEFA3
MPO
LCN2
LTF
MMP8
CAMP
d
cell_fine
DEFA3
MPO
LCN2
LTF
MMP8
CAMP

Supplementary Figure S11. Validation of developing neutrophil population in severe COVID-19 patients from datasets other than Wilk et al. 2020.

a) UMAP embedding showing the plasma cell and epithelial cell (RACK1) subpopulations in the severe COVID-19 patients of Zhang et al. 2020 dataset (n=1320 cells)
b) UMAP embedding showing the plasma cell and epithelial cell (RACK1) subpopulations in the severe COVID-19 patients of Lee et al. 2020 dataset (n=245 cells)
c) UMAP embedding showing the plasma cell and epithelial cell (RACK1) subpopulations in the severe COVID-19 patients of Liao et al. 2020 dataset (n=830 cells)
d) UMAP embedding showing the plasma cell and epithelial cell (RACK1) subpopulations in the severe COVID-19 patients of Chua et al. 2020 dataset (n=1884 cells)

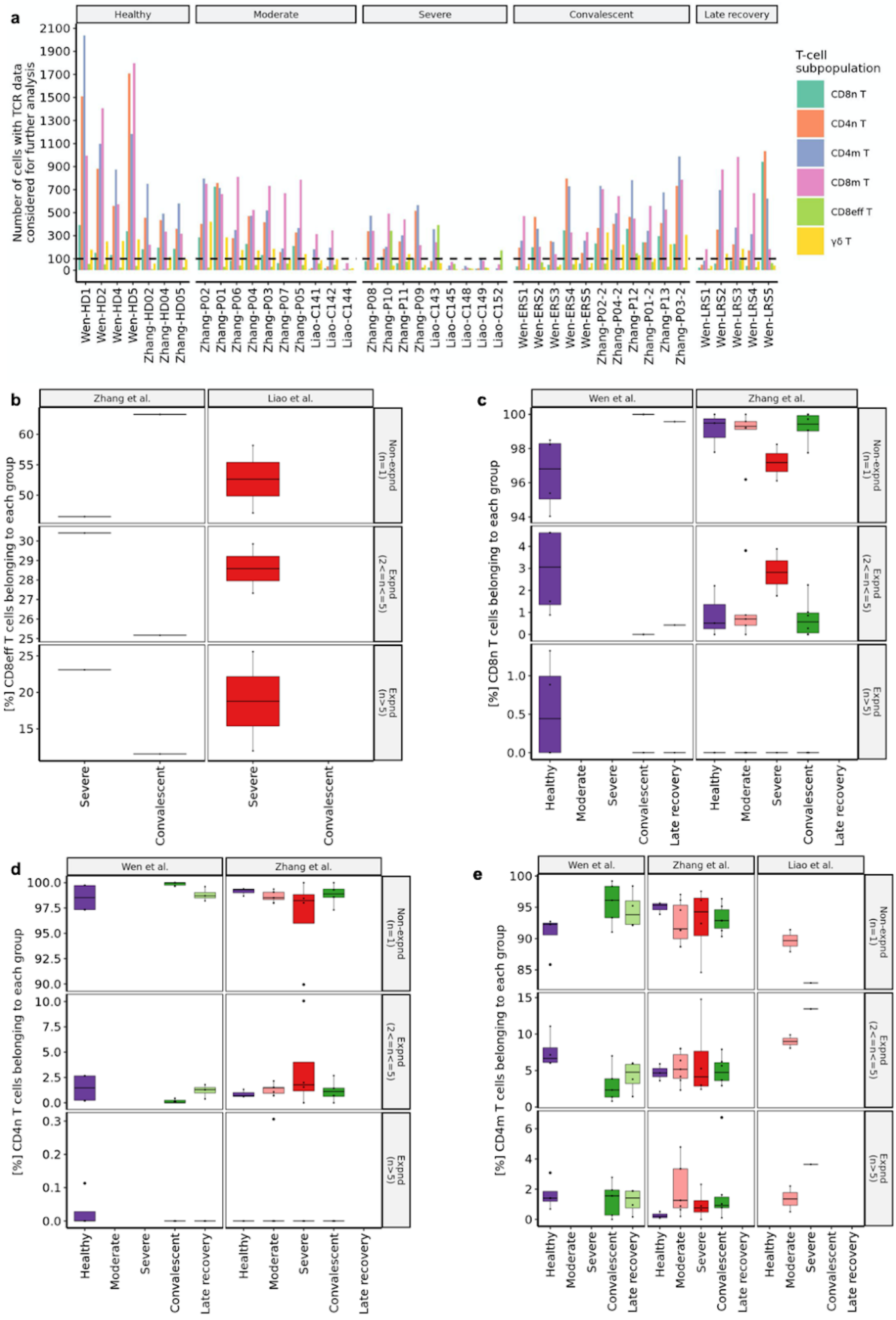
a
Healthy
Moderate
Severe
Convalescent
Late recovery
Number of cells with TCR data considered for further analysis
T-cell subpopulation
CD8n T
CD4n T
CD4m T
CD8m T
CD8eff T
γδ T
Wen-HD1
Wen-HD2
Wen-HD4
Wen-HD5
Zhang-HD02
Zhang-HD04
Zhang-HD05
Zhang-P02
Zhang-P01
Zhang-P06
Zhang-P04
Zhang-P03
Zhang-P07
Zhang-P05
Liao-C141
Liao-C142
Liao-C144
Zhang-P08
Zhang-P10
Zhang-P11
Zhang-P09
Liao-C143
Liao-C145
Liao-C148
Liao-C149
Liao-C152
Wen-ERS1
Wen-ERS2
Wen-ERS3
Wen-ERS4
Wen-ERS5
Zhang-P02-2
Zhang-P04-2
Zhang-P12
Zhang-P01-2
Zhang-P13
Zhang-P03-2
Wen-LRS1
Wen-LRS2
Wen-LRS3
Wen-LRS4
Wen-LRS5
b
Zhang et al.
Liao et al.
[%] CD8eff T cells belonging to each group
Non-expnd (n=1)
Expnd (2<=n<=5)
Expnd (n>5)
Severe
Convalescent
c
Wen et al.
Zhang et al.
[%] CD8n T cells belonging to each group
Healthy
Moderate
Severe
Convalescent
Late recovery
d
Wen et al.
Zhang et al.
[%] CD4n T cells belonging to each group
e
Wen et al.
Zhang et al.
Liao et al.
[%] CD4m T cells belonging to each group

Supplementary Figure S12. Limitations of current TCR data include insufficient number of effector $CD8^+$ T-cells in most of the samples and less than 100 T-cells each in the samples derived from BALF (Liao et al. 2020)

a) Bar plots showing the total number of cells belonging to each T-cell subpopulation in each sample that had the corresponding TCR data, had at least one productive heavy chain and at least one productive light chain. Here, samples with prefix “Wen" belong to the study Wen et al. 2020, samples with prefix “Zhang” belong to the study Zhang et al. 2020 and samples with prefix “Liao” belong to the study Liao et al. 2020.

b) **to e)** Box plots comparing the percentage of different T-cells subpopulations belonging to non-expanded clonotypes (Non-expnd (n=1)), expanded clonotypes with greater than 2 and less than 5 clones (Expnd (2<=n<=5)) and expanded clonotypes with greater than 5 clones (Expnd (n>5)) at each stage across four different studies. **b)** Effector $CD8^+$ T-cells **c)** naive $CD8^+$ T-cells **d)** naive $CD4^+$ T-cells **e)** memory $CD4^+$ T-cells. Colors denote the stage of the patient. All differences were analyzed using two-sided unpaired Wilcoxon rank sum tests with Bonferroni correction and p-values <0.05 are reported.

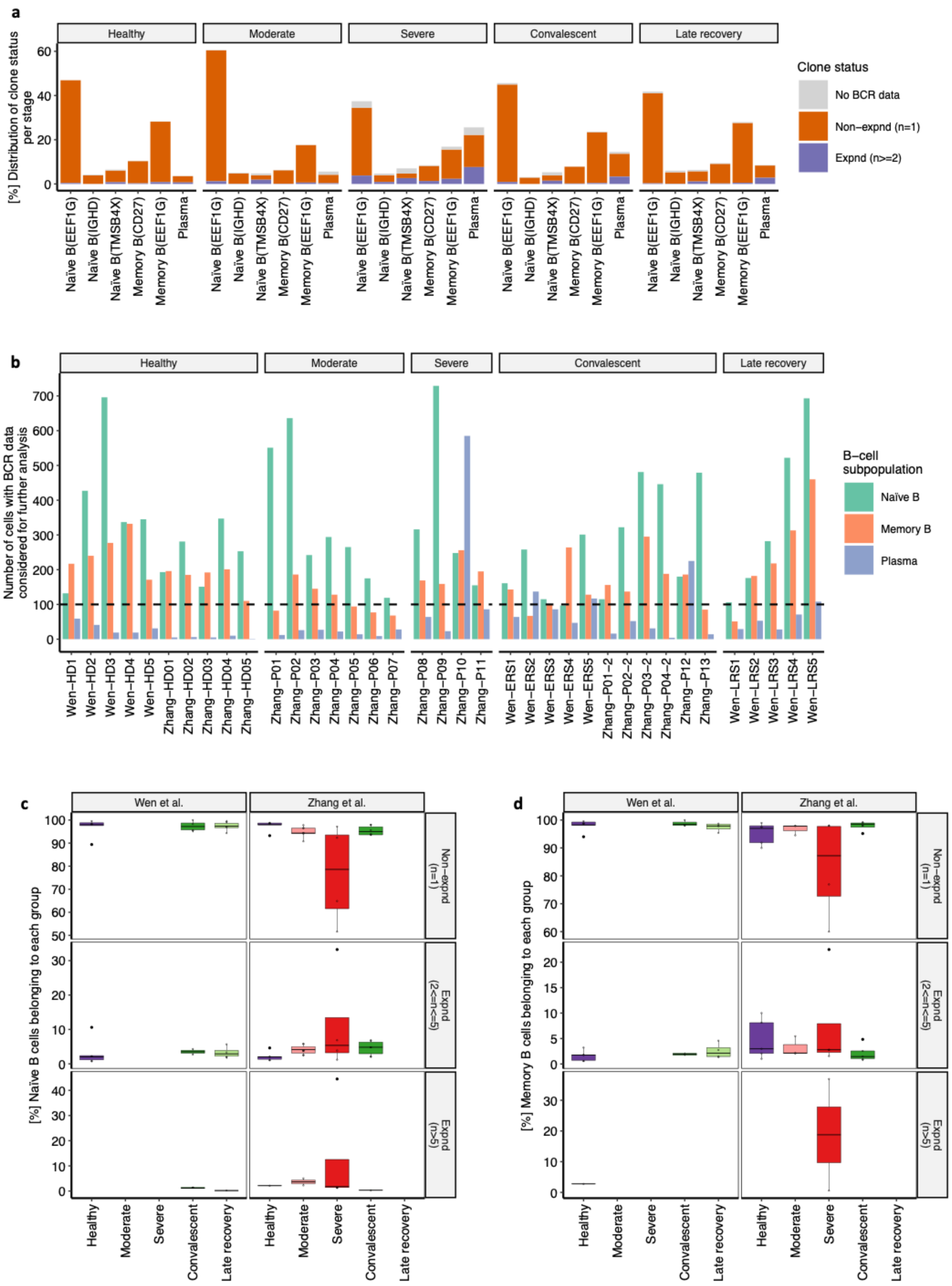


Supplementary Figure S13. Limitations of current BCR data include insufficient number of Plasma cells and memory B-cells in most of the samples along with presence of study-specific stages.

**a)** Bar plots showing the percentage of cells at each stage belonging to the specific B-cell subpopulation as well as clone status. Here, grey color denotes no BCR data, orange color denotes non-expanded clone status (Non-expnd) and blue color denotes expanded clone status (Expnd).

**b)** Bar plots showing the total number of cells belonging to each B-cell subpopulation in each sample that had the corresponding BCR data, had at least one productive heavy chain and at least one productive light chain. Here, samples with prefix “Wen" belong to the study Wen et al. 2020 and samples with prefix “Zhang” belong to the study Zhang et al. 2020.

**c)** **to d)** Box plots comparing the percentage of **c)** naïve B-cells and **d)** memory B-cell subpopulations belonging to non-expanded clonotypes (Non-expnd (n=1)), expanded clonotypes with greater than 2 and less than 5 clones (Expnd (2<=n<=5)) and expanded clonotypes with greater than 5 clones (Expnd (n>5)) at each stage across four different studies. Colors denote the stage of the patient. All differences were analyzed using two-sided unpaired Wilcoxon rank sum tests with Bonferroni correction and p-values <0.05 are reported.

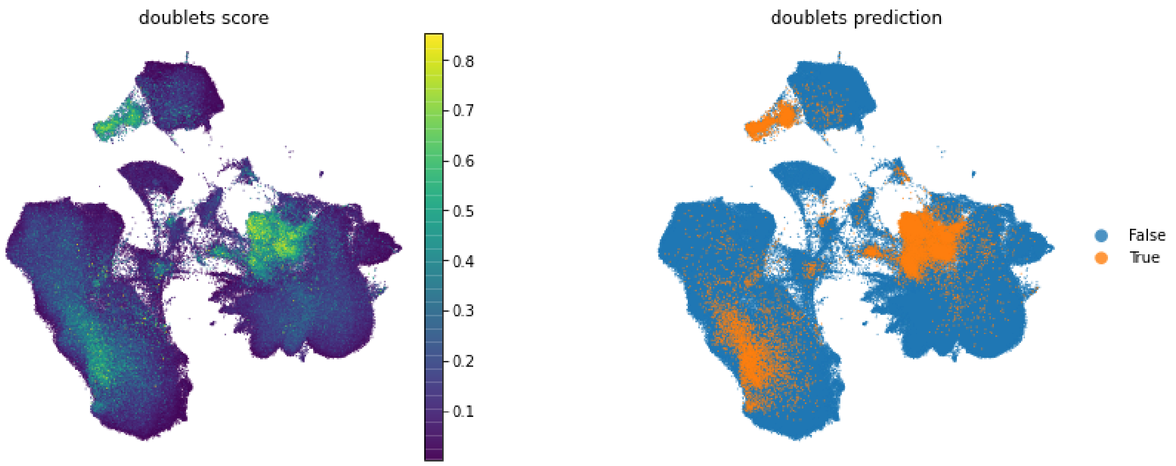


Fig S14. Doublets prediction results from Scrublet.

### Supplementary Notes

#### Interactive visualization on cellxgene through usegalaxy.eu

The first time that you do this, you will need to go through all the steps. On subsequent visits you can just visit the link that you obtain on step 6. The interactive visualisation link will expire after 1 month possibly, but a new one can be generated again from step 4 and on.

If you don’t have an account at usegalaxy.eu, register for one at https://usegalaxy.eu/login and then login with that account.

Once logged:

1.- Go to https://usegalaxy.eu/u/pmoreno/h/garg-et-al-2020-covid-19-single-cell-immune-response-meta-analysis find the Galaxy history with the data.

2.- Press the import button (+) at the upper right corner next to “About this History”:

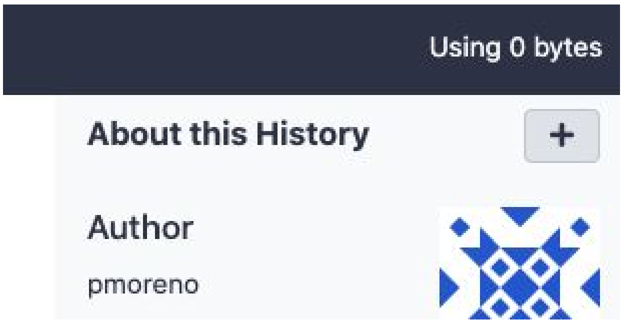


3.- Press Import in the dialog

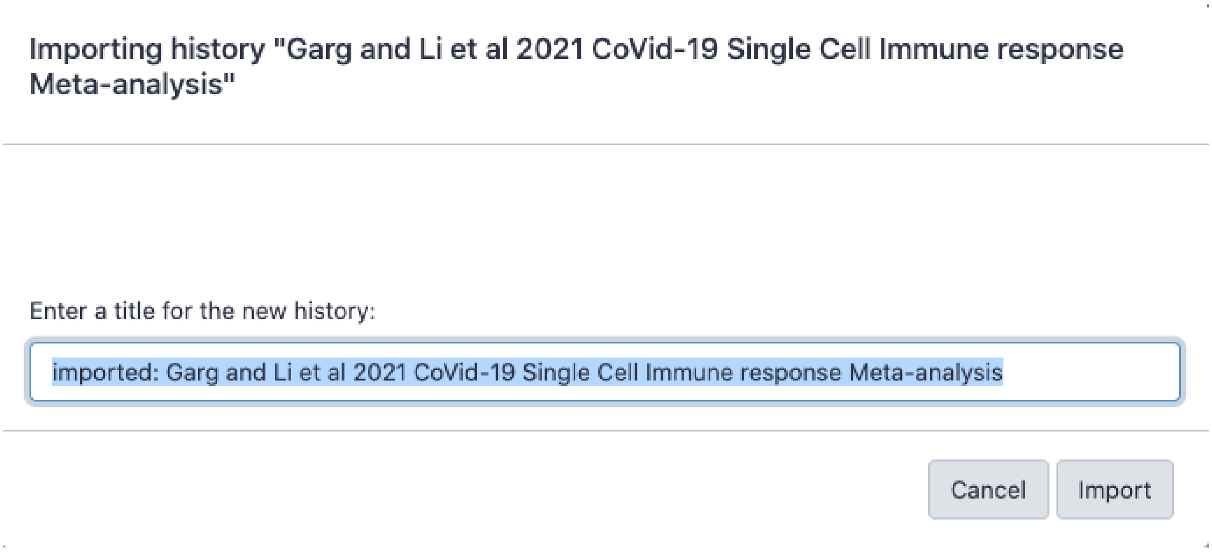


4.- Go to https://usegalaxy.eu/root?tool_id=interactive_tool_cellxgene

5.- Making sure that the field Concatenate Dataset that appear in the middle has this dataset

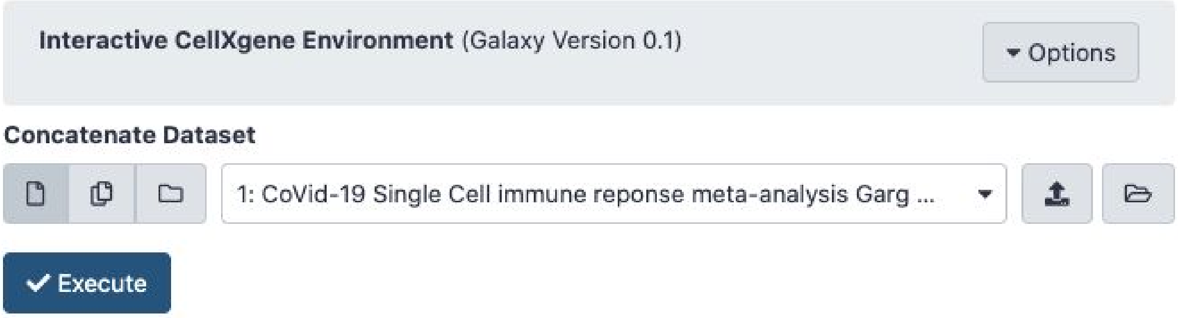


Click on Execute, you should see execution to the right on the history.

6.- The central page will show this at the top:

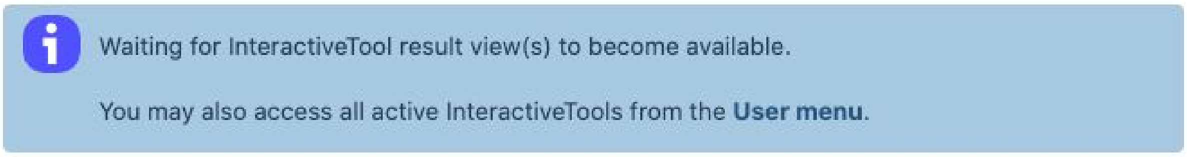


And then after a short while (this can take longer if the cluster is under heavy use) it should change to this:

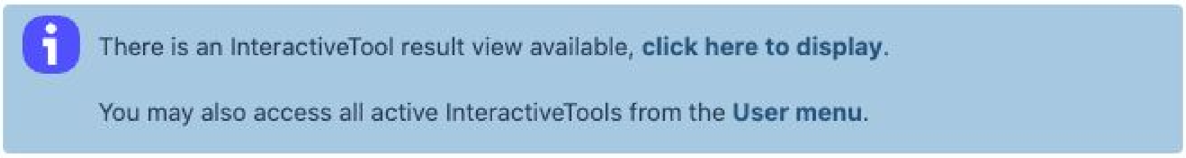


Click on "click here to display" and you should arrive to cellxgene with the merged dataset loaded. This link should be valid for at least 30 days, keep it so that you can go by directly to your cellxgene instance next time (if you have lost the link, there are indications to get it back after the final step).

7.- Enter a name on the text field to create an annotation collation to start exploring the dataset on cellxgene:

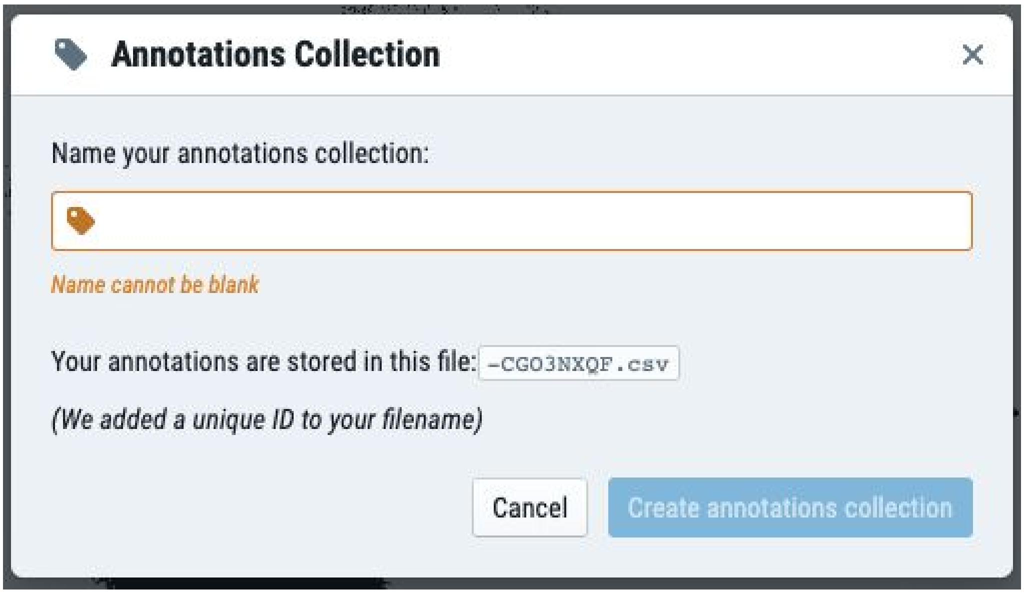


What if I didn't keep the link to the cellxgene instance?

1.- Once you are logged in, go to the Active InteractiveTools item in the User menu.

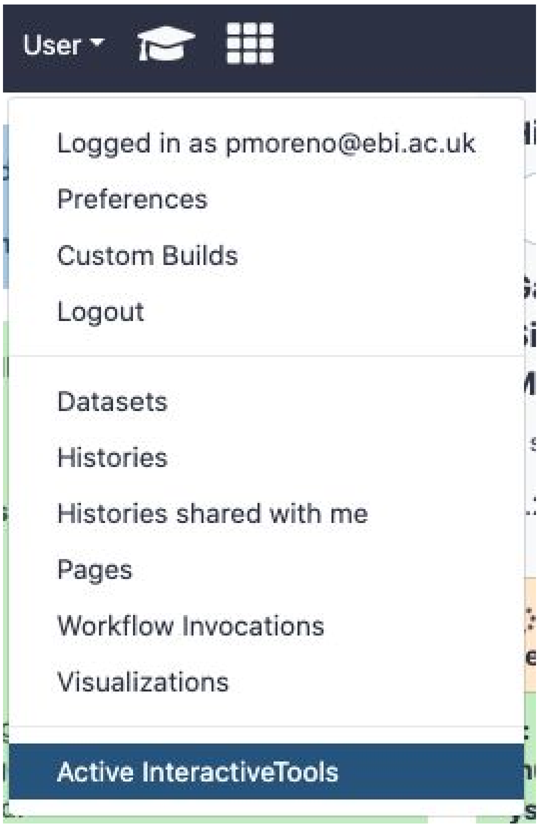


2.- You should see an entry like this, click on the name:

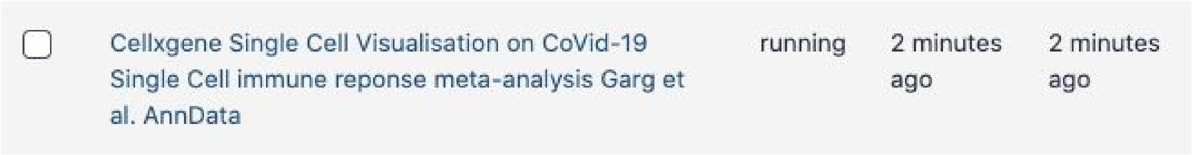


Clicking on it should take you to your own deployment of cellxgene with the dataset.
